# An atlas of expressed transcripts in the prenatal and postnatal human cortex

**DOI:** 10.1101/2024.05.24.595768

**Authors:** Rosemary A. Bamford, Szi Kay Leung, V. Kartik Chundru, Aaron R. Jeffries, Jonathan P. Davies, Alice Franklin, Xinmu Chen, Andrew McQuillin, Nicholas Bass, APEX consortium, Emma Walker, Paul O’Neill, Ehsan Pishva, Emma L. Dempster, Eilis Hannon, Caroline F. Wright, Jonathan Mill

**Affiliations:** Department of Clinical and Biomedical Sciences, University of Exeter Medical School, University of Exeter, Barrack Road, Exeter, UK; Department of Biosciences, University of Exeter, Exeter, UK; Molecular Psychiatry Laboratory, Windeyer Institute of Medical Sciences, Research Department of Mental Health Sciences, University College London, London, UK; Maastricht University, Maastricht, Netherlands

## Abstract

Alternative splicing is a post-transcriptional mechanism that increases the diversity of expressed transcripts and plays an important role in regulating gene expression in the developing central nervous system. We used long-read transcriptome sequencing to characterise the structure and abundance of full-length transcripts in the human cortex from donors aged 6 weeks post-conception to 83 years old. We identified thousands of novel transcripts, with dramatic differences in the diversity of expressed transcripts between prenatal and postnatal cortex. A large proportion of these previously uncharacterised transcripts have high coding potential, with corresponding peptides detected in proteomic data. Novel putative coding sequences are highly conserved and overlap *de novo* mutations in genes linked with neurodevelopmental disorders in individuals with relevant clinical phenotypes. Our findings underscore the potential of novel coding sequences to harbor clinically relevant variants, offering new insights into the genetic architecture of human disease. Our cortical transcript annotations are available as a resource to the research community via an online database.

## MAIN

Alternative splicing (AS) is a post-transcriptional regulatory mechanism that acts to generate distinct mRNA molecules from a single pre-mRNA molecule. It plays an important role in development of the vertebrate central nervous system^1^ with altered RNA splicing being implicated in numerous neurodevelopmental disorders^2,3^. Long-read transcript sequencing has facilitated the unambiguous characterization of full-length isoforms^4,5^, enabling the identification of novel transcript structures and coding regions that potentially harbour genetic variants predisposing to disease. In this study we used Oxford Nanopore Technologies (ONT) sequencing to generate a comprehensive atlas of transcripts expressed in the human cortex. We identify thousands of novel transcripts, with changes in isoform abundance across development and between males and females. We show that many of these novel coding sequences are highly conserved and overlap *de novo* variants in genes linked with neurodevelopmental disorders. Our findings underscore the potential of novel coding sequences to harbor clinically relevant variants, offering new insights into the genetic architecture of human disease.

### Characterizing full-length transcripts in the cortex using long-read sequencing

We generated long-read whole transcriptome sequencing data from human cortex tissue dissected from 47 donors (21 male and 26 female, 26 prenatal and 21 postnatal) aged 6 weeks post-conception (WPC) to 83 years (**Fig. 1a** and **Supplementary Table 1**). In total, we generated 207.71 million (mean = 4.42 million per sample) high-quality ONT reads with a total yield of transcriptome sequencing data across all samples of 368.88 Gb (**Fig. 1b** and **Supplementary Table 2**). Raw data were pre-processed using a bespoke pipeline and filtered to remove single-exon intergenic transcripts and single-exon transcripts from multi-exonic genes that likely reflect intra-priming events or nascent RNA^6^ (**Methods**). Reads in our final dataset corresponded to 1,895,360 unique transcripts that mapped to 36,149 annotated genic features (16,269 coding genes, 11,922 non-coding RNA genes, 3,200 pseudogenes, 3,835 fusion genes and 923 antisense genes) with a mean transcript length of 1.87 kb (SD = 1.13 kb, mean number of exons = 7.38). Rarefaction sensitivity curves confirmed that we had sequenced to saturation and that the final filtered dataset was representative of the true cortical transcript population **(Extended Data** Fig. 1). Comparison with a previous long-read human cortex transcriptome dataset generated by our group on tissue from a small number of donors^4^ highlighted that the majority (n = 56,252, 86%) of previously detected transcripts were also detected in the current analysis (**Extended Data** Fig. 2). Compared to these overlapping transcripts, the 2,285,616 additional transcripts identified in the current study were characterised by significantly lower expression (mean read counts: common transcripts = 348, transcripts identified in current study = 51; Mann-Whitney P < 2 x 10^-16^). There was considerable diversity in the number of unique transcripts identified per gene (mean = 52.43, SD = 110.33), with half of all annotated genic features (n = 18,106, 50.1%) found to express more than unique isoforms (**Fig. 1c**). *HNRNPK*, encoding a ubiquitously expressed RNA binding protein that interacts with heterogeneous nuclear RNA, was characterized by the largest number of unique transcripts (n = 3,051 isoforms) in the cortex (**Extended Data** Fig. 3 and **Supplementary Table 3**). Our finalized transcript annotations and downstream expression findings are available as an open resource to the research community via a searchable online database available from our laboratory website (https://www.epigenomicslab.com/brainisoform/).

**Figure 1.**
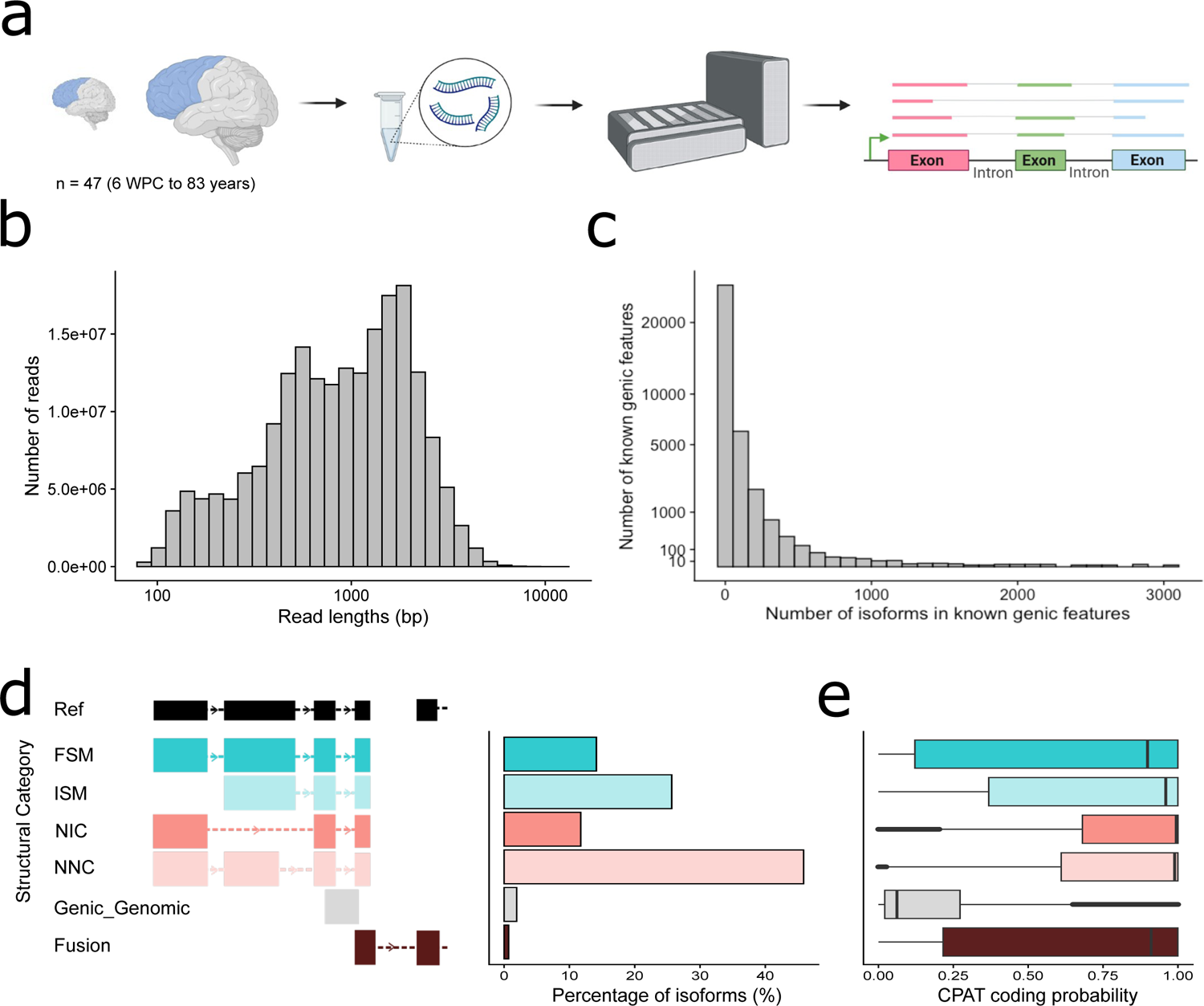
Long-read sequencing identifies novel transcripts of annotated genic features in the human cortex. **a)** An overview of the experimental approach used to generate ONT long-read whole transcriptome data on human cortex tissue from 47 donors aged 6 wpc to 83 years old. **b)** The distribution of full-length transcript reads after basecalling, alignment and filtering (see **Methods**). **c)** The number of unique transcripts identified for each known genic feature detected in the human cortex. **d)** The proportion of transcripts split by structural category. A transcript was classified as being a ‘full splice match’ (FSM) if it aligned with the GENCODE v44 (hg38) reference genome with the same internal splice junctions and contained the same number of exons, an ‘incomplete splice match’ (ISM) if it had fewer external exons than the reference transcript, ‘novel in catalogue’ (NIC) if it represented a novel transcript containing a combination of known donor or acceptor sites, and ‘novel not in catalogue’ (NNC) if it represented a novel transcript with at least one novel donor or acceptor site. ‘Genic’ transcripts partially overlap exons in a known gene and ‘fusion’ transcripts incorporate transcribed sequences from two or more adjacent genes. **e)** The predicted coding probability of transcripts split by structural category highlighting the high coding potential of novel transcripts. Coding probability was determined using CPAT^9^ which characterises ORFs among detected transcripts to predict coding potential.

### A large proportion of expressed transcripts are novel compared with annotated genic features

A notable proportion of transcripts expressed from annotated genic features were not present in existing GENCODE v44 (hg38) annotations. In total we identified 1,091,522 ‘novel’ transcripts (57.6% of all transcripts annotated to known genes) associated with 21,960 (60.7%) annotated genic features (mean transcript length = 1.96 kb, SD = 1.08 kb, mean number of exons = 8.58). The majority of these transcripts were classified by SQANTI3^7^ as ‘novel not in catalog’ (NNC) (n = 869,221, 79.6% of all novel transcripts) and contained at least one novel donor or acceptor site (**Fig. 1d**). The remaining novel transcripts were primarily ‘novel in catalog’ (NIC) (n = 222,301, 20.3% of all novel transcripts) comprising either new combinations of already annotated splice junctions or novel splice junctions formed from already annotated donors and acceptors. As a group, novel transcripts were significantly less abundant than known transcripts (Mann-Whitney P < 2 x 10^-308^). Full-splice match (FSM) transcripts were the most abundant transcript classification for the majority (53.9%) of annotated genic features; among 16,199 known protein-coding genes detected in the cortex, FSM transcripts were most abundant for 9,830 (60.7%) and of these 7,656 (77.9%) corresponded to the MANE Select transcript^8^. Despite this, for 11,694 (32.3%) of annotated genic features (including 1,824 (11.3%) protein-coding genes with at least one observed transcript) we did not detect a FSM transcript expressed in the cortex, with novel transcripts being the most highly expressed isoform for 21.8% of genic features (18.4% within protein-coding genes). The ribosomal protein gene *RPS27A* had the two most abundantly expressed novel transcripts (ONT2_3223_23931 and ONT2_3223_23974 (both NNC)); together these account for ∼92% of all transcript reads from this gene (**Extended Data** Fig. 4) with both isoforms characterised by the same shorter predicted coding sequence compared to the GENCODE v44 reference transcript. We assessed the coding probability of transcripts using the Coding-Potential Assessment Tool (CPAT)^9^ which characterises ORFs among detected transcripts to predict coding potential (see **Methods**) (**Fig. 1e**). Overall we identified a high level of coding potential with 1,459,651 (62.3%) detected transcripts predicted to be protein coding, including a high proportion of novel transcripts (NIC =81.4%, NNC = 80.2%).

### Considerable transcript diversity in the human cortex across development

AS is an important mechanism regulating gene expression in the developing central nervous system^10^. We explored differences in transcript expression across cortex development by comparing whole transcriptome data generated on cortex tissue from prenatal (n = 26 (11 male and 15 female), mean age = 25.6 weeks post-conception (WPC), range = 6-28 WPC,) and postnatal (all samples > 1 year old, n = 19 (10 male and 9 female), mean age = 34.3 years, range = 2-83 years) donors (**Supplementary Table 1**). There was no significant difference in the number of full-length reads generated between these groups (prenatal cortex: 4.49 M mean processed reads, postnatal cortex: 4.33 M mean processed reads, t-test P = 0.85, **Supplementary Table 2**). 2,433 known genic features (6.73% of all detected) only had reads in prenatal cortex (including 309 protein-coding genes), and 2,214 (6.12% of all detected) only had reads in postnatal cortex (including 441 protein-coding genes) (**Supplementary Table 3**). Among the remaining 31,502 known genic features (87.1% of those detected) present in both prenatal and postnatal cortex, many were characterised by striking differences in transcript diversity across development (**Supplementary Table 3**).

*DLGAP5* had the greatest relative enrichment of prenatally-expressed transcripts (prenatal cortex = 143 transcripts, postnatal cortex = 1 transcript, **Fig. 2a**), whilst *DLGAP1-AS4* had the greatest enrichment of postnatally-expressed transcripts (postnatal cortex = 148 transcripts, prenatal cortex = 1 transcript) (**Supplementary Table 3**). Although many changes in transcript diversity across development parallel overall gene expression differences between prenatal and postnatal cortex (**Supplementary Tables 3-4**), many genes showed dramatic changes in the number of distinct transcripts without any difference in gene-level expression. The overall expression of the 5-Hydroxytryptamine Receptor 4 gene (*HTR4*), for example, was not significantly different between prenatal and postnatal cortex but there were nine distinct transcripts in the postnatal cortex compared to two in the postnatal cortex (**Supplementary Table 3**). Although the majority of transcripts annotated to known genic features were detected in both the prenatal and postnatal cortex (n = 1,184,402 shared transcripts, 75.5% of all prenatal detected transcripts, 78.3% of all postnatal detected transcripts), a notable proportion were unique to either the prenatal (n = 383,335; 20% of all transcripts) or postnatal (n = 327,623; 17% of all transcripts) cortex (**Extended Data** Fig. 5**, Supplementary Table 3**). Of note, the majority of novel transcripts were detected in both prenatal and postnatal cortex (**Extended Data** Fig. 6). The most abundant prenatal-only transcript (ONT1_5283_13707, 2,842 full length reads) was an FSM isoform of *NHLH1*, a basic helix-loop-helix transcription factor involved in the development of the nervous system^11^, whereas the most abundant postnatal-only cortex transcript (ONT18_5132_2320, 8,026 full length reads) was an isoform of *MBP*, a highly isomorphic gene encoding a protein involved in the myelination of oligodendrocytes and Schwann cells in the developing nervous system^12^ (**Supplementary Table 3**).

**Figure 2.**
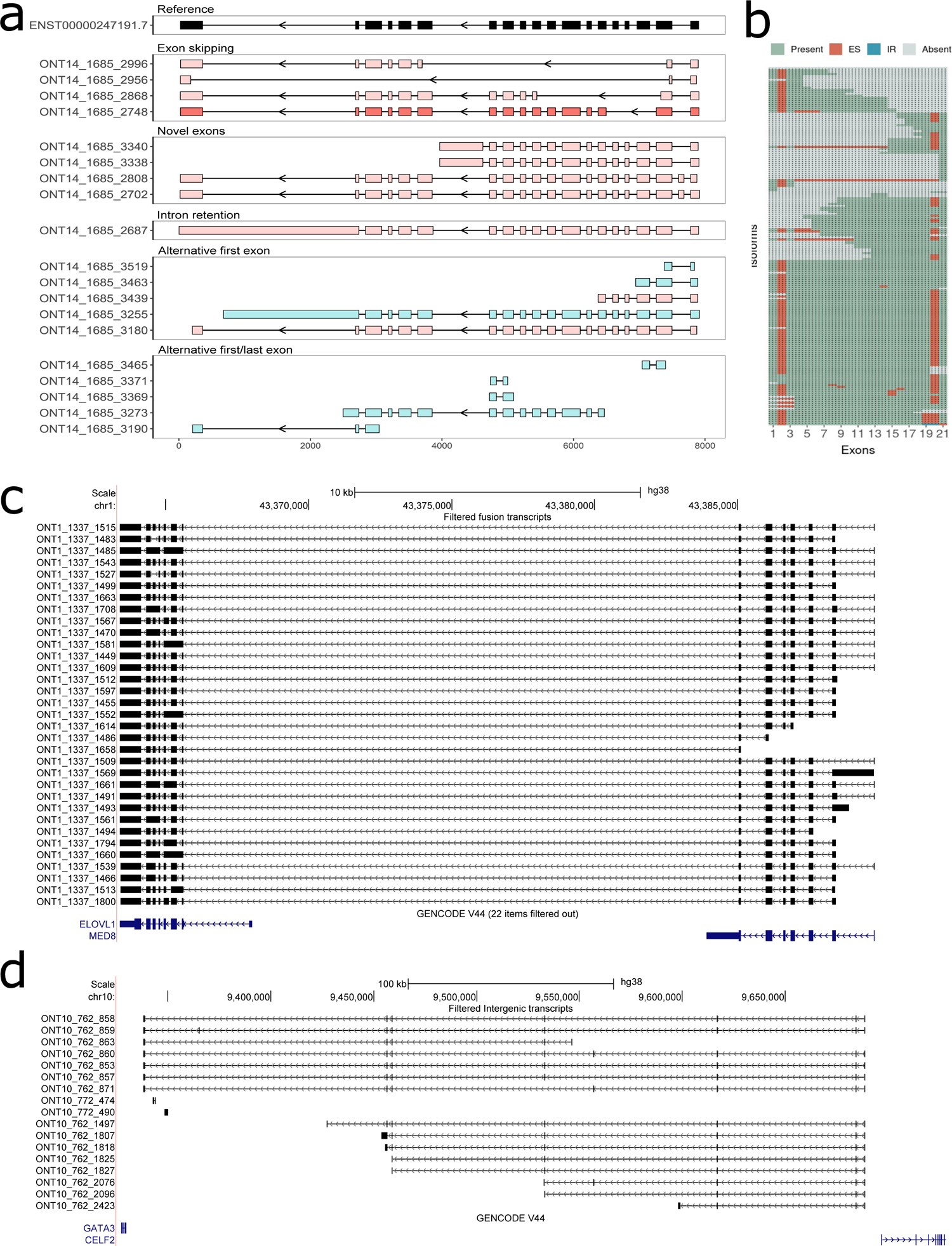
Long-read transcript sequencing identifies widespread alternative splicing events, fusion transcripts and putative novel genes in the human cortex. **a)** Examples of novel *DLGAP5* isoforms highlighting different alternative splicing events (exon skipping, novel exons, intron retention, alternative first exon and alternative first/last exon) occurring in the prenatal cortex. Transcripts are coloured according to structural category (FSM = turquoise; ISM = light turquoise; NIC = dark pink; NNC = light pink; genic = gray; fusion = brown). **b)** A cluster dendrogram depicting the transcriptional landscape of *DLGAP5*. Each row corresponds to a distinct isoform and each column represents a distinct exon. **c)** Fusion transcripts incorporating transcribed sequences from *ELOVL1* and *MED8*. Each represents a putative coding fusion transcript that includes sequence from both genes. A full list of fusion transcripts can be found in **Supplementary Table 6. d)** A putative novel multi-exonic genic feature located between *GATA3* and *CELF3* on chromosome 10. The transcripts ONT10_762_860 and ONT10_762_859 have 8 exons. A full list of putative novel coding genes can be found in **Supplementary Table 7.**

### Alternative splicing events make a major contribution to transcriptional diversity in the cortex

AS is known to be a major driver of transcriptional diversity in the central nervous system^13^. To fully describe AS events for individual genes, we developed *FICLE*, a bespoke analysis pipeline that compares splice junctions between long-read-derived and reference transcripts, enabling a gene-level overview of all detected isoforms^14^. We used *FICLE* to characterise the frequency of multiple different AS events (exon skipping (ES), alternative first exon use (AF), alternative last exon use (AL), differential A3’ and A5’ splice site use, and intron retention (IR)) associated with transcripts expressed from all multi-exonic protein-coding genes in the human cortex (**Fig. 2b**, **Supplementary Table 5**). Overall ES and A5A3 were the most prevalent AS events in the human cortex (**Extended Data** Fig. 7) although there were differences in the prevalence of specific AS events across development, with a significantly higher frequency of ES (t-test: P < 2.2 x 19^-16^) and IR (t-test: P = 1.03 x 10^-3^) in the prenatal cortex. Visualisation of all alternative splicing events associated with protein-coding genes expressed in the human cortex is available in our online resource (https://www.epigenomicslab.com/brainisoform/).

### Identification of fusion transcripts and novel multi-exonic transcripts not associated with annotated genes

13,204 detected transcripts (0.5% of total) involving 5,227 (14.5%) annotated genic features were classified as ‘fusion transcripts’, incorporating transcribed sequences from two or more adjacent genes resulting from transcriptional readthrough^15^. We further filtered these transcripts to retain those incorporating >1 exon from each gene, resulting in 4,650 robust fusions incorporating sequence from 2,243 annotated genic features, including 3,570 transcripts representing fusions of two protein-coding genes (**Supplementary Table 6**).

Although often considered to reflect transcriptional ‘noise’ with limited functional relevance^16^, we used CPAT to show that 86% of these transcripts were predicted to be protein-coding (CPAT coding probability > 0.364). Overall, fusion transcripts were significantly more abundant in the postnatal cortex compared to the prenatal cortex (after adjusting for total read count, t =-2.8, df = 6,353, P = 0.005). The most abundant fusion transcript, expressed in both prenatal and postnatal cortex, incorporates multiple exons from *FKBP1A* and *SDCBP2* located adjacently on chromosome 20. Interestingly, we find evidence of fusion transcripts incorporating sequences from genes that have been previously reported as being highly coexpressed (e.g. *ELOVL1* and *MED8* on chromosome 1^17^, **Fig. 2c**). In addition to fusion transcripts, we identified 32,892 transcripts not assigned to a previously annotated genic feature (**Supplementary Table 7**). We further filtered these to exclude antisense transcripts overlapping known genes, leaving 8,675 transcripts mapping to 4,460 putative ‘novel genes’. A number of these putative novel genes were found to be highly expressed, and overall novel gene transcripts were more abundant in the postnatal cortex (after adjusting for total read counts, t-test t=-2.6, df=11597, P = 0.009) (**Fig. 2d**). Although the vast majority of these transcripts were predicted to be non-coding, 331 (4%) were predicted to be protein-coding (CPAT coding probability>0.364, **Supplementary Table 7**).

### Human cortex proteomics data provides evidence for the translation of novel transcripts

We generated mass spectrometry (MS)-based proteomics data on an independent set of postnatal human cortex samples (n = 35, 18 male and 17 female aged 61 to 101 years, **Supplementary Table 8**) to assess whether novel (NIC and NNC) transcripts were translated by searching for ORFs predicted from CPAT (see **Methods**). We detected 779,792 peptides mapping to 276,200 transcripts in our cortex dataset, finding evidence for the translation of 229,594 novel transcripts in the postnatal cortex including evidence supporting the translation of novel coding sequences resulting from different AS events. For example, peptides assigned to novel transcripts of *STMN2* - a gene robustly associated with Alzheimer’s disease^18^ - provide evidence for translation of protein isoforms with exon skipping and alternative splice sites (**Fig. 3a**), and peptides assigned to a novel transcript of *SHC1* confirm the translation of a novel coding exon (**Fig. 3b**). A full list of novel transcripts confirmed from novel peptides using human cortex mass spectrometry data is provided in Supplementary Table 9.

**Figure 3.**
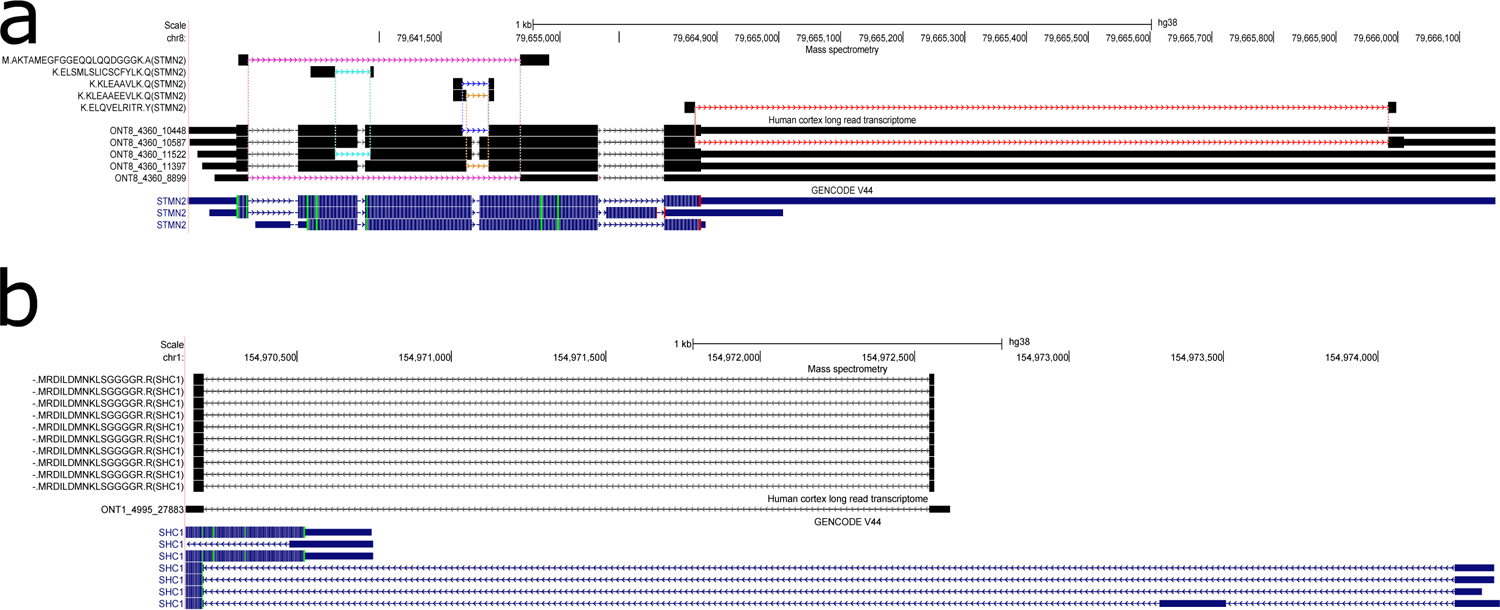
Evidence that novel transcripts are translated into novel peptides. **a)** Novel transcripts of *STMN2* characterised by exon skipping and the use of alternative splice sites. Human cortex mass spectrometry proteomic data identifies novel peptides validating these alternative splicing events (colours indicate transcripts and their corresponding peptides detected by mass spectrometry). **b)** A transcript of *SHC1* containing a novel coding exon that differs strikingly from the canonical start site. Mass spectrometry protein data confirms this is translated into a novel peptide in the postnatal cortex.

### Differential transcript expression and use between prenatal and postnatal cortex

We next sought to identify specific differentially expressed transcripts (DETs) between prenatal and postnatal cortex, focusing on a subset of transcripts (n = 497,873) characterised by a minimum of 10 reads across all samples (see **Methods**). We identified 65,864 DETs (FDR < 0.05) annotated to 15,069 genic features (**Supplementary Table 10** and **Fig. 4a**) with the top-ranked DET being a known isoform of *MBP*, which was dramatically up-regulated in postnatal cortex (ONT18_5132_2313, log2FC = 14.8, FDR = 1.32 x 10^-148^) (**Extended Data** Fig. 8). Overall, there was a significant enrichment of DETs becoming upregulated in the postnatal cortex (n = 39,518 (60%), binomial test P = 9.88 x 10^-324^). A large proportion of DETs were novel transcripts (n = 30,292 (46%)) with the top-ranked novel DET also being an isoform of *MBP* (ONT18_5132_2319, log2FC =10.7, SE = 0.56, FDR = 8.83 x 10^-70^) (**Fig. 4b**).

**Figure 4.**
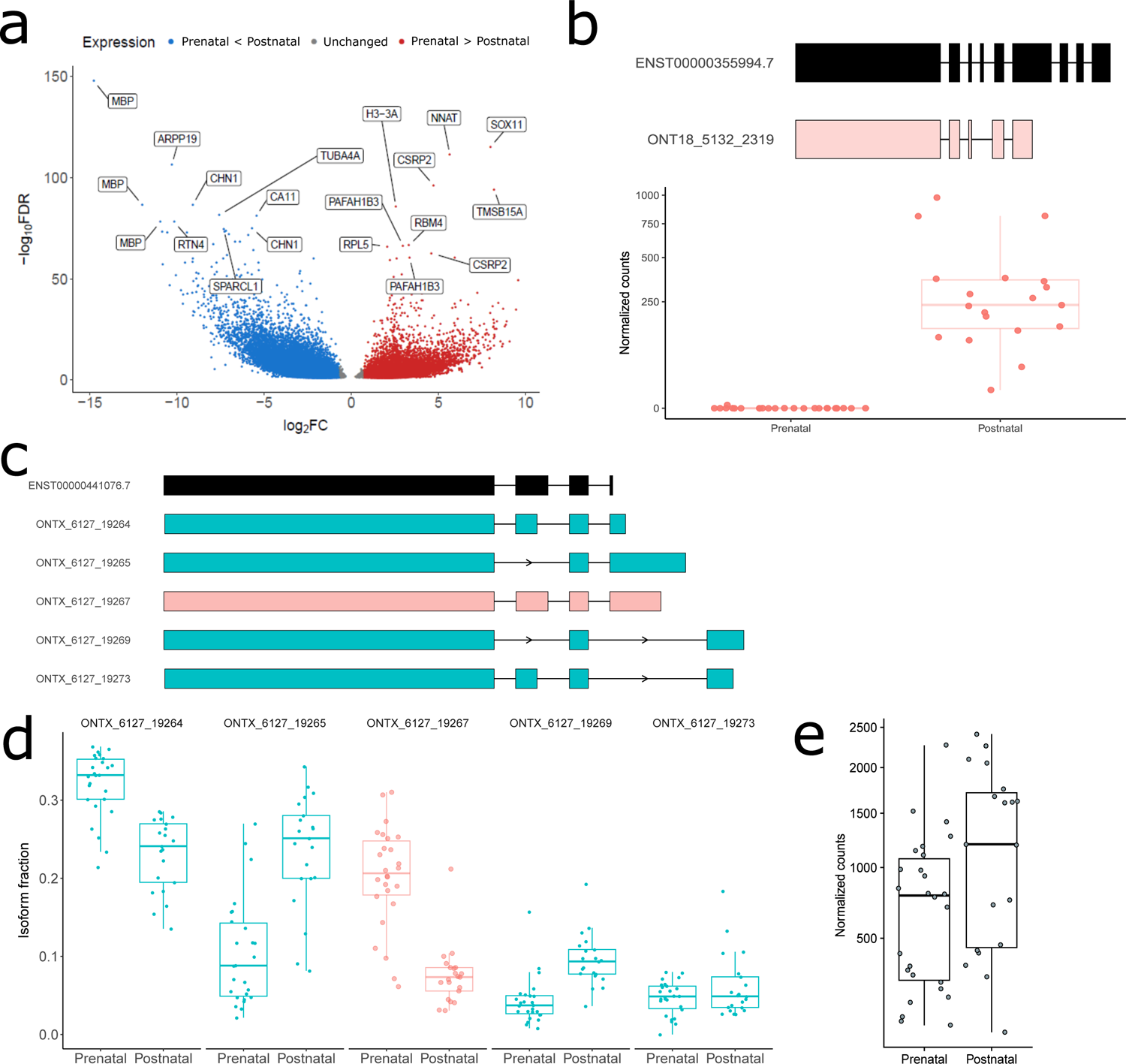
Differential transcript expression and usage between prenatal and postnatal cortex. **a)** Volcano plot highlighting the 65,864 differentially expressed transcripts (FDR < 0.05) between prenatal and postnatal cortex. Gene annotations are shown for the top 20 up-and down-regulated DETs. **b)** The top-ranked novel DET was annotated to *MBP* and characterised by significantly higher expression in postnatal cortex (ONT18_5132_2319, log2FC = 10.7, FDR = 8.83 x 10^-70^). **c)** Tracks depicting the top five most abundant transcripts of *MORF4L2* which is characterised by differential transcript use between prenatal and postnatal cortex (FDR = 5.00 x 10^-89^). **d)** The relative isoform fraction for the five most abundant transcripts of *MORF4L2*. The relative proportion of 4 of these transcripts changed significantly between prenatal and postnatal cortex (ONTX_6127_19264: prenatal cortex (0.31), postnatal cortex (0.23), P = 8.36 x 10^-7^; ONTX_6127_19265: prenatal cortex (0.11), postnatal cortex (0.24), P = 3.95 x 10^-8^; ONTX_6127_19267: prenatal cortex (0.20), postnatal cortex (0.07), P = 7.44 x 10^-12^; ONTX_6127_19269: prenatal cortex (0.04), postnatal cortex (0.10), P = 9.63 x 10^-07^). There was a switch in major *MORF4L2* transcript between prenatal and postnatal cortex, with ONTX_6127_19264 being most abundant in prenatal cortex and ONTX_6127_19265 being most abundant in postnatal cortex. **e)** There was no overall difference in *MORF4L2* gene expression between prenatal and postnatal cortex. Tracks are coloured according to structural category (FSM = turquoise; ISM = light turquoise; NIC = dark pink; NNC = light pink; genic = gray; fusion = brown).

We identified a relatively high number of antisense DETs (n = 7,969 (12.1%)), an interesting observation given their known role in gene regulation and development^19^. In addition to changes in the abundance of specific transcripts we also found evidence for extensive differential transcript use (DTU) - i.e. changes in the relative proportion of different transcripts - between prenatal and postnatal cortex. In total we observed significant DTU at 2,923 genes (**Supplementary Table 11**), with 690 (23.6%) characterised by a switch in the dominant expressed isoform between prenatal and postnatal cortex (**Extended Data** Fig. 9) and 1,029 (35.20%) of these changes occurring without an overall gene-level expression difference between prenatal and postnatal cortex. The top-ranked DTU gene between prenatal and postnatal cortex was *MORF4L2*, which was characterised by dramatic changes in the abundance of specific transcripts across development despite no overall difference in gene expression (**Fig. 4c-e**).

### Sex differences in transcript expression in the developing cortex

Given evidence for sex-biassed gene expression^20^, we explored the extent to which specific transcripts were differentially expressed in the cortex between males and females. We identified 65 sex DETs (FDR < 0.05) annotated to 27 genes (**Extended Data** Fig. 10, **Supplementary Table 12**); as expected the majority (n = 57 (87.7%)) of these were transcripts of genes located on the sex chromosomes (**Extended Data** Figure 11). The top-ranked autosomal sex DET was a known transcript of the *ADD3* gene on chromosome 10, encoding a membrane-cytoskeleton-associated protein, which was significantly up-regulated in cortex tissue from male donors compared to female donors (ONT10_4920_1919, log2FC = −3.74, FDR = 7.78 x 10^-4^) (**Extended Data** Fig. 12). This transcript was also one of eight sex DETs that was also characterised by differential expression between prenatal and postnatal cortex (**Extended Data** Fig. 13). The top-ranked sex DTU gene was the imprinted gene *GNAS* (FDR = 1.32 x 10^-256^) (**Extended Data** Fig. 14, **Supplementary Table 13**) which was characterised by a different dominant isoform in male and female cortex but no overall gene-level sex difference in expression (**Extended Data** Fig. 14, **Supplementary Table 14**).

### Differences in untranslated region (UTR) length is a major driver of transcript diversity between pre- and post-natal cortex

We used an established long-read proteogenomic pipeline^21^ to further explore the coding potential of all 2,341,868 unique transcripts (**Methods**) finding that 1,405,075 (60%) were predicted to be protein-coding (CPAT coding probability > 0.364). Of these, 590,904 (42%) were predicted to have a distinct ORF resulting in 567,945 unique protein isoforms after filtering (**Methods**). There was a small but significant difference in predicted coding sequence (CDS) length between prenatal and postnatal cortex (mean = 837.21bp, and 839.75bp respectively, t-test p=4.7 x 10^-116^); however, we found a much larger difference in lengths of untranslated regions (UTRs), with a second shorter peak in prenatal samples (mean = 645.03bp (prenatal) vs 802.62bp (postnatal), t-test p <10^-324^) (**Extended Data** Fig. 15). This difference is driven primarily by variation in 3’UTR length (mean = 474.18bp (prenatal) vs 610.79bp (postnatal), t-test p <10^-324^), but also paralleled in 5’UTRs (mean = 171.85bp (prenatal) vs 192.82bp (postnatal), t-test p <10^-324^). Comparing transcripts containing the same ORF and predicted to translate to the same peptide sequence, we observed that the transcriptional diversity was driven exclusively by variation in the 5’ and 3’UTR. This is an interesting observation given the importance of 3’UTRs in neural function and development^4^ and their hypothesized role in the aetiology of neurodevelopmental disorders^22^.

### Ultradeep targeted sequencing reveals additional rare isoforms expressed from disease-associated genes

Given the evidence linking splicing dysfunction to neurodevelopmental disorders^23^ we extended our whole transcriptome data using ultradeep targeted sequencing (see **Methods**) to perform an unprecedented characterisation of transcripts expressed from 241 genes robustly associated with autism (133 high-confidence ASD genes from the SFARI gene database^24^), schizophrenia (32 genes identified via exome-sequencing by the SCHEMA consortium^25^ and 71 genes prioritised by the most recent GWAS analysis^26^), and severe developmental disorders (142 genes in the DDG2P list of curated developmental disorder genes^27^) (**Supplementary Table 15**) profiling cortex tissue from a subset of donors (24 prenatal cortex and 16 postnatal cortex, **Supplementary Table 1**). Across these genes we validated 82% of the transcripts identified using whole transcriptome profiling, including 79% of the identified novel transcripts (**Extended Data** Fig. 16 and **Supplementary Table 16**). As expected, the targeted approach yielded substantially more (60,611) transcripts across these genes, including 44,390 additional novel transcripts not included in existing GENCODEv44 annotations. Together these data provide unprecedented insight into the transcriptional landscape of genes implicated in neurodevelopmental disorders with widespread alternative splicing (**Fig. 5**) and a large number of novel coding transcripts.

**Figure 5.**
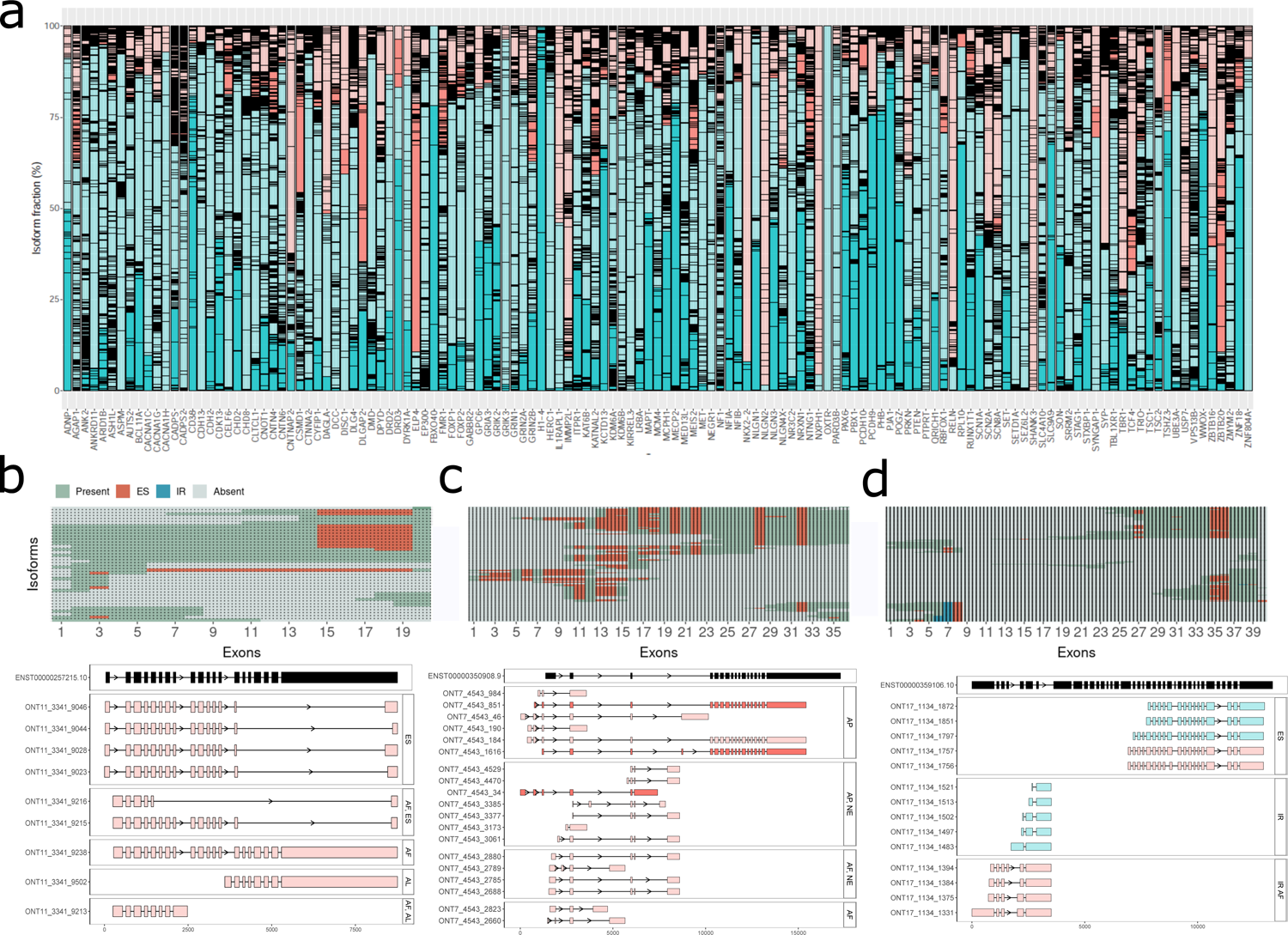
Transcript diversity for genes associated with neurodevelopmental disorders. **a)** Detected isoforms and their usage for 134 high-confidence autism genes from the SFARI gene database^24^ using combined whole transcriptome and targeted long-read sequencing. This list of autism genes includes 18 overlapping schizophrenia^25^ and 41 monoallelic DDG2P genes^27^. The bars are coloured according to structural category (FSM = turquoise; ISM = light turquoise; NIC = dark pink; NNC = light pink; genic = gray; fusion = brown). The lines across the bars delineate individual isoforms. The isoform fraction per gene is determined by dividing the mean normalized read counts of the associated isoforms across all the samples, by the total normalized mean read counts of all the associated isoforms. Cluster dendrograms depicting the transcript landscape of **b)** *DAGLA*, a schizophrenia (SCHEMA) gene, **c)** *FOXP2,* a category 1 SFARI gene and a strong monoallelic DDD gene, **d)** *CACNA1G,* a category 2 SFARI gene, SCHEMA gene and a strong monoallelic DDD gene. Tracks for transcripts of **e)** *DAGLA*, **f)** *FOXP2* and **g)** *CACNA1G*.

### Novel coding sequences are enriched for highly-conserved bases

In total we identified 2.35Mb of novel predicted CDS - i.e. regions of the genome not previously assigned as coding in a GENCODEv44 transcript - contained within 31,869 transcripts annotated to 7,252 known protein-coding genes (**Supplementary Table 17**). These regions were found to be highly conserved with a significantly higher mean phyloP (cactus241way^28^) conservation score (0.70) than both GENCODE UTR sequences (mean phyloP score = 0.50, t-test P < 2×10^-308^) and random intronic regions (mean phyloP score = 0.11, t-test P < 2×10^-308^). The proportion of highly conserved bases (phyloP>2.27) in these novel CDS (0.14) was also significantly higher than reference GENCODE UTRs (0.11, 2-sample Z test P < 2×10^-30^) and random intronic regions (0.03, 2-sample Z test P < 2×10^-308^) (**Fig 6a**).

**Figure 6.**
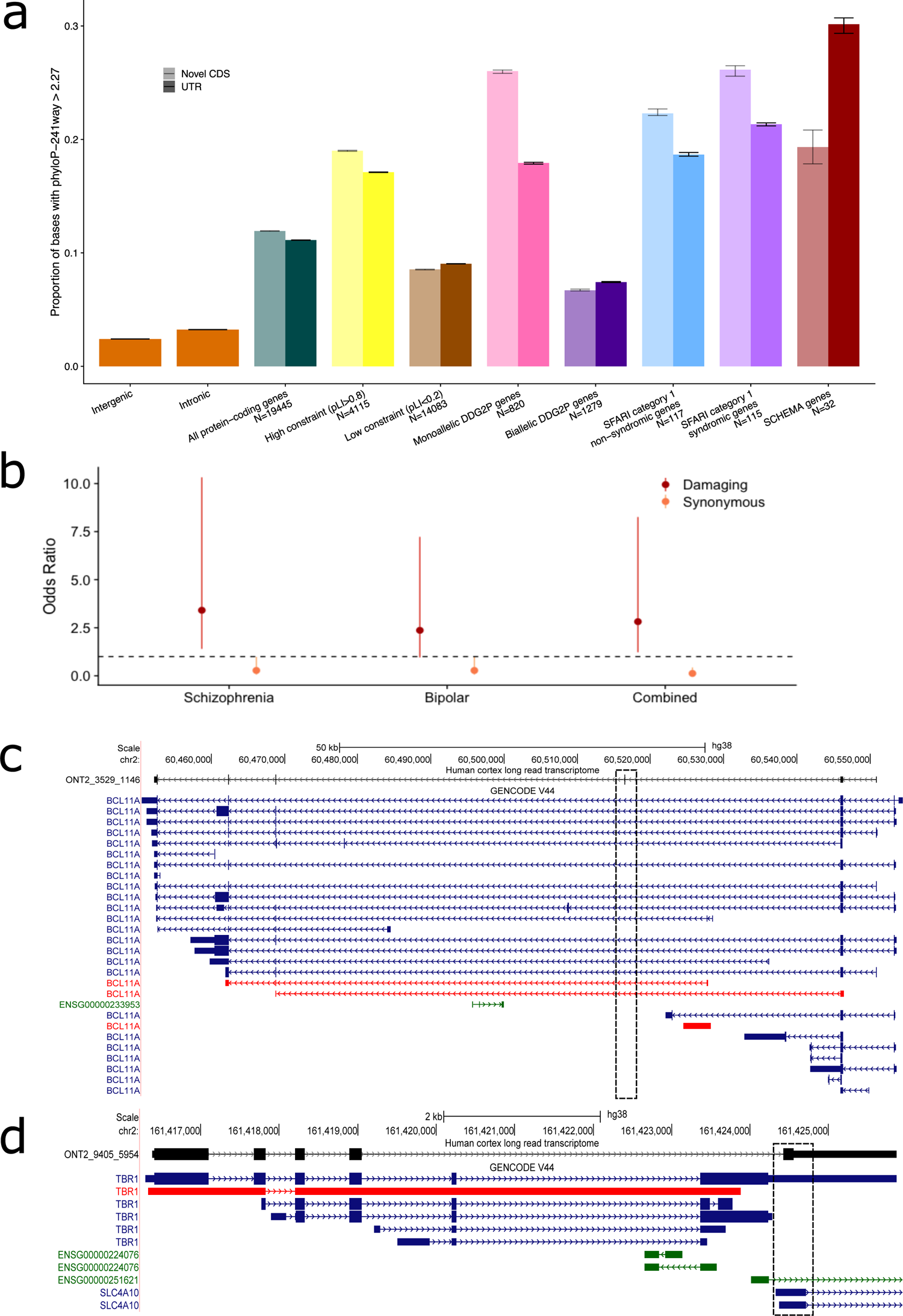
Novel coding sequences harbour pathogenic variants associated with neurodevelopmental disorders. **a)** The proportion of highly conserved bases (PhyloP>2.27) in random intronic sequences, random intergenic sequences, and novel coding sequences and GENCODEv44 UTRs in all genes, high and low constraint genes, and genes associated with neurodevelopmental disorders. **b)** There is a significant enrichment of rare (gnomADv2 MAF<1×10^-4^) non-synonymous (‘damaging’) variants in schizophrenia and bipolar disorder patients (OR = 2.8, P=0.02) compared to controls in whole exome sequencing data. **c)** A novel predicted coding *BCL11A* transcript (ONT2_3529_1146) containing a novel exon. Shown is a UCSC track of the transcript alongside existing GENCODE V44 annotation. The putative pathogenic stop gained *de novo* mutation occurs in the novel exon shown (see **Table 1**). **d)** A novel predicted coding TBR1 transcript (ONT2_9405_5954) containing novel CDS in the 3’UTR. Shown is a UCSC track of the transcript alongside existing GENCODE V44 annotation. The putative pathogenic stop gained *de novo* mutation occurs in the last exon within the novel CDS.

**TABLE 1.**
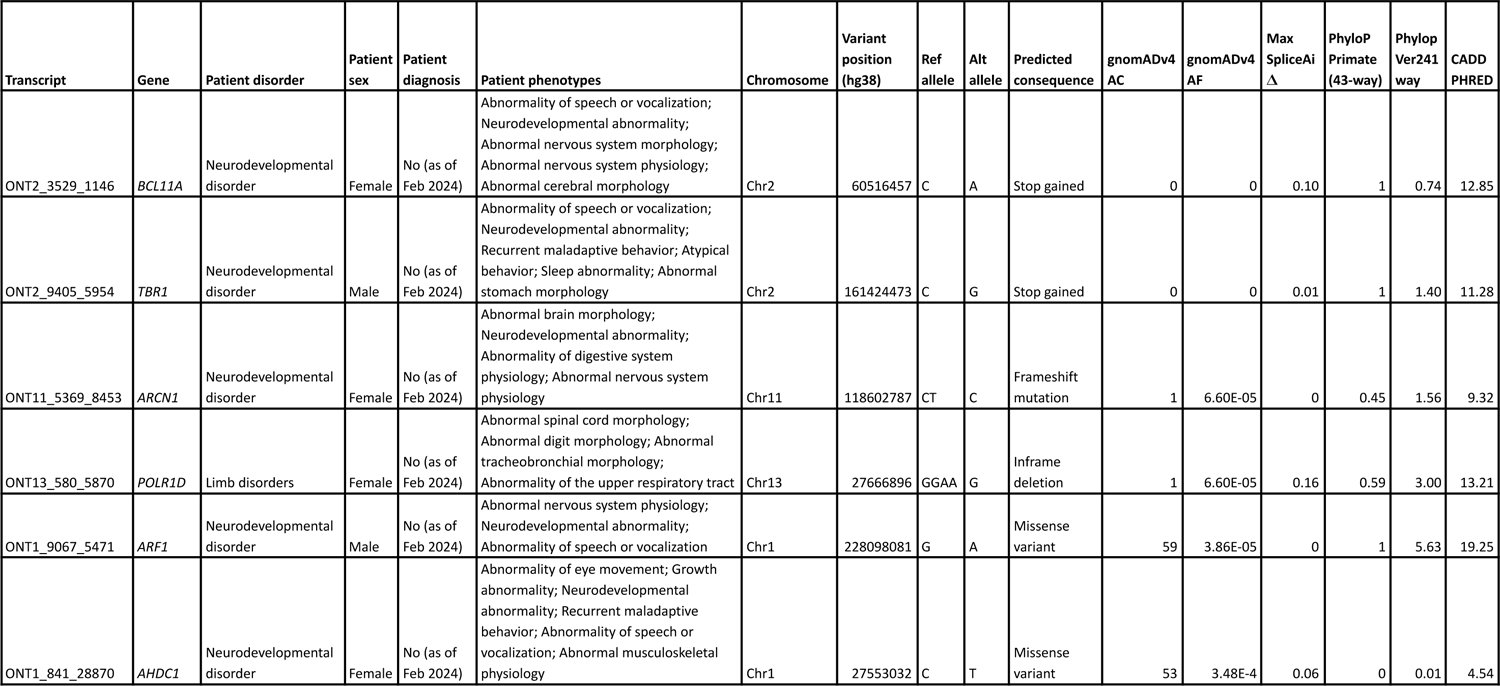
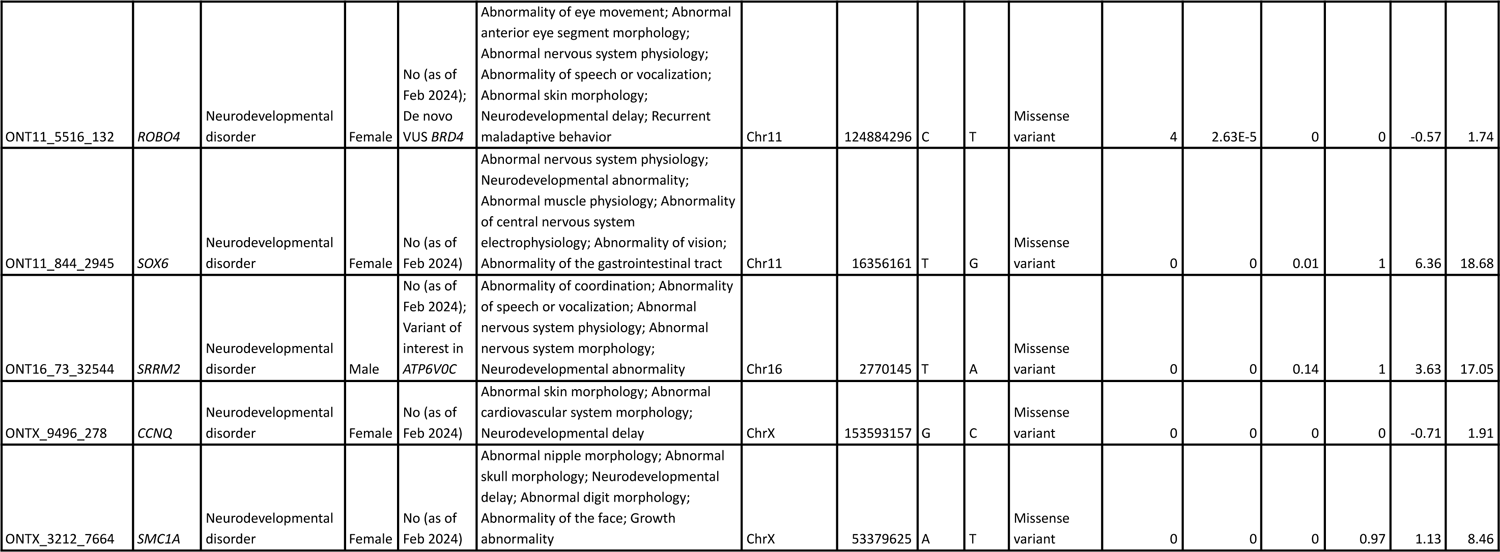
Novel coding sequences harbour putative non-synonymous *de novo* variants in genes associated with neurodevelopmental disorders. We used stringently filtered *de novo* mutations identified in neurodevelopmental disorder cases from the Genomics England 100,000 Genomes Project cohort (n = 6,702 trios)^32^, identifying 11 previously undetected *de novo* non-synonymous mutations overlapping novel CDS amongst monoallelic DDG2P^33^ genes.

This enrichment was stronger in highly constrained genes (pLI>0.8) and novel transcribed sequences mapping to unconstrained genes (pLI<0.2)^29^ showed a depletion of highly conserved bases compared to GENCODE UTRs (**Figure 6a**). Amongst genes associated with neurodevelopmental phenotypes, this enrichment was particularly strong in the monoallelic (dominant) DDG2P genes and SFARI autism genes (**Fig. 6a)**.

### Novel coding sequences harbour rare candidate diagnostic *de novo* variants in genes associated with neurodevelopmental disorders

The novel putative coding regions identified in disease-associated genes have implications for the identification of causal variants outside of the canonical coding regions that may be missed using approaches such as exome sequencing. Hypothesising that novel CDS might harbour previously undetected pathogenic variants, we integrated our novel transcript annotations with sequencing data from patients with neurodevelopmental phenotypes. First we used published exome sequencing data^30^ from psychosis patients (n = 1,551 schizophrenia and 1,949 bipolar disorder) and 1,117 controls^25,31^ (see **Methods**). Although these data are limited by the coverage of exome sequencing which does not capture any novel exons, there was an overall significant enrichment of rare (gnomADv2 MAF<1×10^-4^) non-synonymous damaging variants in novel CDS captured by exome-seq in patients (OR = 2.8, P = 0.02, **Fig. 6b**) highlighting the utility of our novel transcript annotations for burden testing. We next extended our analyses to focus on whole genome sequencing (WGS) data to look for putative diagnostic *de novo* mutations in highly penetrant monogenic neurodevelopmental disorders. We used stringently filtered *de novo* mutations identified in neurodevelopmental disorder cases from WGS data from the Genomics England 100,000 Genomes Project cohort (n = 6,702 trios)^32^, identifying 17 previously undetected *de novo* mutations overlapping novel CDS amongst monoallelic DDG2P^33^ genes. We re-annotated these variants, using the Ensembl Variant Effect Predictor (VEP)^34^ to predict the consequence of the variants in the novel transcripts (see **Methods**); 11 (64%) were predicted to be non-synonymous (corresponding to three predicted loss-of-function (pLoF), and eight missense or inframe indel variants) in previously undiagnosed probands, with the other six being synonymous variants (**Table 1**). Of particular interest was a stop-gained variant in *BCL11A* located in a novel exon identified in a transcript (ONT2_3529_1146) not present in GENCODE (**Fig. 6c**). This variant was not present in the gnomADv4 database^35^ and has no predicted splicing effect according to SpliceAI^36^. The proband with this *de novo* mutation had no tier 1 or tier 2 variants prioritised by Genomics England^37^, and their clinical phenotypes closely matched those of known pathogenic *BCL11A* mutations including intellectual disability, hypotonia, microcephaly, behaviour problems, abnormal brain structure and seizures^38,39^. We also detected a stop-gained variant in *TBR1* located in a novel putative coding exon in a transcript not present in GENCODE (ONT2_9405_5954); this variant is also not present in gnomADv4 and has no predicted splicing effect (**Fig. 6d**). While there is little known about the TBR1 protein, other pathogenic ClinVar variants have been detected in the final exon, and the end of the protein is predicted to have a protein-binding function^40^. The phenotype of this individual aligns closely with *TBR1-*related disorder, including intellectual disability, speech delays, autistic traits, aggressive behaviour, movement disorders, hypotonia, and fine motor delay^41^, suggesting a likely causal role for this novel variant. Our findings underscore the potential of novel coding sequences to harbor clinically relevant variants, offering new insights into the genetic architecture of neurodevelopmental disorders.

## CONCLUSION

In this study we leveraged long-read sequencing to characterise the structure and abundance of full-length transcripts in the human cortex. Our study revealed thousands of novel transcripts and uncovered dramatic differences in the diversity of expressed transcripts between prenatal and postnatal cortex. A large proportion of these previously uncharacterised transcripts have high coding potential, with corresponding peptides detected in proteomic data generated on human cortex tissue. Novel putative coding sequences were highly conserved suggesting that genetic variants in these regions may have important pathogenic consequences with implications for genetic burden testing and existing exome-sequencing studies that are focused on known coding regions. By integrating the novel coding sequences identified in this study with whole genome sequencing data from the Genomics England 100,000 Genomes Project we identified *de novo* mutations in genes linked with neurodevelopmental disorders in individuals with relevant clinical phenotypes. Taken together, our findings expand understanding about the complexity of gene expression in the human cortex and underscore the importance of alternative splicing in shaping transcript diversity in the central nervous system. Our comprehensive atlas of cortical transcripts serves as a valuable resource for the scientific community, paving the way for further exploration into the molecular underpinnings of brain development and disease.

## ACKNOWLEDGEMENTS

This work was supported by a grants from the Simons Foundation for Autism Research (SFARI) (grant number 573312, awarded to J.M. and grant number 809383, awarded to the APEX consortium), a grant from the UK Medical Research Council (grant MR/R005176/1, awarded to J.M.), and Wellcome Trust [226083/Z/22/Z]. The human prenatal material was provided by the Human Developmental Biology Resource (funded by MRC/Wellcome Trust grant (099175/ Z/12/Z) (https://www.hdbr.org). Sequencing infrastructure was supported by a Wellcome Trust Multi User Equipment Award (WT101650MA, awarded to J.M.) and Medical Research Council (MRC) Clinical Infrastructure Funding (MR/M008924/1, awarded to J.M.). This research was made possible through access to data in the National Genomic Research Library, which is managed by Genomics England Limited (a wholly owned company of the Department of Health and Social Care). The National Genomic Research Library holds data provided by patients and collected by the NHS as part of their care and data collected as part of their participation in research. The National Genomic Research Library is funded by the National Institute for Health Research and NHS England. The Wellcome Trust, Cancer Research UK and the Medical Research Council have also funded research infrastructure.

This study was supported by the National Institute for Health and Care Research Exeter Biomedical Research Centre. The views expressed are those of the author(s) and not necessarily those of the NIHR or the Department of Health and Social Care. Schizophrenia, bipolar disorder and control sample exome sequencing data was supported by the Medical Research Council (grant G1000708), the Stanley Center for Psychiatric Research at the Broad Institute, the Stanley Medical Research Institute and University College London Hospitals NHS Foundation Trust NIHR BRC. We acknowledge the supply of samples from the Quebec Suicide Brain Bank at Douglas Mental Health University Institute, Canada; The Mount Sinai NBTR (NIH Brain and Tissue Repository); Human Developmental Biology Resource at the IGM, Newcastle upon Tyne; The Guy’s and St Thomas’ Research Biobanks; The Institute of Child Health, University College London, London; Kings College London in association with the MRC London Brain Bank for Neurodegenerative Diseases and funders of the Brain Bank; The South London and Maudsley NHS Foundation Trust; The London Neurodegenerative Diseases Brain Bank, Brains for Dementia Research; JJ Peters VA Medical Center; the Oxford Brain Bank, supported by the Medical Research Council (MRC), the NIHR Oxford Biomedical Research Centre and the Brains for Dementia Research programme, jointly funded by Alzheimer’s Research UK and Alzheimer’s Society; The Neuropathology Brain Bank at the University of Pittsburgh, School of Medicine Department of Psychiatry; The Stanley Medical Research Institute Brain Collection courtesy of Drs. Michael B. Knable, E. Fuller Torrey, Maree J. Webster, and Robert H. Yolken; The Human Brain and Spinal Fluid Resource Center (HBSFRC) NIH Neurobiobank, UCLA. Postnatal human cortex brain samples for mass spectrometry were provided by the Brains for Dementia Research (BDR) cohort. BDR is jointly funded by Alzheimer’s Research UK (ARUK) and the Alzheimer’s Society in association with the UK Medical Research Council.

## AUTHOR CONTRIBUTIONS

JM designed and supervised the study. JM, EH, ELD, CFW and the APEX consortium obtained funding. RAB led and supervised laboratory experiments with help from JPD. RAB undertook primary data pre-processing with analytical and computational input from EW ARJ, PO and AF. RAB, SKL, VKC led the analysis of genetic variant data with input from CFW, XC, NB and AM. SKL designed and generated the interactive database. SKL and EP generated mass spectrometry data on human cortex tissue. RAB, SKL, VKC and JM interpreted the data and drafted the manuscript. All authors reviewed the final manuscript before submission.

## AUTHOR INFORMATION

**APEX Consortium:** Dwaipayan Adhya, Carrie Allison, Bonnie Auyeung, Rosemary Bamford, Simon Baron-Cohen, Richard Bethlehem, Tal Biron-Shental, Graham Burton, Jonathan Davies, Dori Floris, Alice Franklin, Lidia Gabis, Daniel Geschwind, David M. Greenberg, Yuanjun Gu, Alexandra Havdahl, Alexander Heazell, Rosemary Holt, Matthew Hurles, Yumnah Khan, Meng-Chuan Lai, Madeline Lancaster, Michael Lombardo, Hilary Martin, Jose Gonzalez Martinez, Jonathan Mill, Mahmoud K. Musa, Kathy Niakan, Adam Pavlinek, Lucia Dutan Polit, Marcin Radecki, David Rowitch, Laura Sichlinger, Deepak Srivastava, Alex Tsompanidis, Florina Uzefovsky, Varun Warrier, Elizabeth Weir, Xinhe Zhang.

## METHODS

### Human cortex tissue samples

Prenatal cortex tissue (n = 26 donors) was obtained from the Human Developmental Biological Resource (HDBR) (https://www.hdbr.org). Ethical approval for the HDBR was granted by the Royal Free Hospital research ethics committee under reference 08/H0712/34 and HTA material storage licence 12220. Post-natal prefrontal cortex tissue from individuals with minimal neuropathology (n = 21 donors) was obtained from registered brain banks in the UK, Canada and US (full details provided in **Supplementary Information** and **Supplementary Table 1**). Briefly, subjects were approached in life for written consent for brain banking, and all tissue donations were collected and stored following legal and ethical guidelines. Demographic data (sex and age) for each donor is detailed in **Supplementary Table 1.** ∼20 mg of tissue was homogenized in Trizol (Thermo Fisher Scientific, UK) and total RNA was isolated using Direct-zol columns (Zymo, USA). RNA samples were quantified and RNA integrity numbers (RIN) derived using the Bioanalyzer High Sensitivity RNA 6000 Pico kit (Agilent, UK).

### Oxford Nanopore Technologies (ONT) whole transcriptome library preparation, sequencing and data pre-processing

Total RNA was converted to cDNA using Maxima H Minus reverse transcriptase (Thermo Fisher Scientific, UK) and amplified with 14 cycles of PCR using LongAmp *Taq* Master Mix (New England Biolabs, UK). cDNA was quantified using the Qubit DNA High sensitivity assay (Invitrogen, UK). ONT library preparation was performed using the PCR-cDNA barcoding kit SQK-PCB109 (Oxford Nanopore Technologies, UK). Libraries were sequenced on the ONT PromethION platform using R9.4.1 FLO-PRO002 flow cells (Oxford Nanopore Technologies, UK) and base-called using *Guppy* (v5.0.12). Resulting *fastq* files were processed through the *Pychopper* pipeline (https://github.com/epi2me-labs/pychopper) to orientate and trim full-length transcript sequences.

### Bespoke barcoding of cDNA and targeted enrichment

Barcodes were added to a subset of cDNA samples (n = 40, **Supplementary Table 1**) using a modified protocol of the NEBNext Single Cell/Low Input cDNA Synthesis & Amplification Module (New England Biolabs, UK) (**Supplementary Methods** and **Supplementary Table 18)** during reverse transcription of RNA, as previously described^1^. The IDT xGen Custom Hyb Panel Design Tool (https://eu.idtdna.com/pages/tools/xgen-hyb-panel-design-tool) was used to design hybridisation probes against 239 target genes, with at least one probe designed for each know exon (**Supplementary Table 19**). We performed an adapted version of the Pacific Biosciences cDNA Capture Using IDT xGen Lockdown Probes protocol (Part Number 101-604-300 Version 01 (June 2018)^2^) on equimolar pooled barcoded cDNA. Samples were quantified using the Qubit DNA High sensitivity assay (Invitrogen, UK) and library preparation was performed using the ONT SQK-LSK110 ligation kit (Oxford Nanopore Technologies, UK). Libraries were sequences as described above.

### Transcriptome annotation and filtering

Long-read transcriptome sequencing data was analysed using a bespoke pipeline adapted by our group (https://github.com/SziKayLeung/LOGen). Briefly, full-length transcripts were mapped to the human reference genome (GENCODE v44 (hg38)) using *minimap2* (https://github.com/lh3/minimap2) and corrected using *TranscriptClean* (https://github.com/mortazavilab/TranscriptClean). Data from all samples were merged and re-aligned using *pbmm2* (https://github.com/PacificBiosciences/pbmm2) as input to collapsing to full-length transcripts using *Iso-Seq collapse* (https://isoseq.how/). Full length read counts from each individual sample were extracted using custom scripts (adapt_cupcake_to_ont.py, demux_cupcake_collapse.py) (https://github.com/SziKayLeung/LOGen). The merged dataset was annotated using *SQANTI3* (https://github.com/ConesaLab/SQANTI3)^3^ in combination with reference gene annotations (GENCODE v44 (hg38)). A transcript was classified as a Full Splice Match (FSM) if it aligned with the reference genome with the same internal splice junctions and contained the same number of exons. Transcripts matching consecutive, but not all, splice junctions of the reference genome were classified as Incomplete Splice Match (ISM). FSM and ISM are both annotated transcripts to known genes. The Novel in Catalog (NIC) label was used to describe a novel transcript containing a combination of known donor or acceptor sites, and the Novel Not in Catalog (NNC) label was used to describe transcripts containing at least one novel donor or acceptor site. *SQANTI3* filtering was performed using the default JSON file, with modification of “min_cov” from 3 to 0 for non-FSM isoforms. We applied further stringent filtering including discarding mono-exonic intergenic transcripts, and mono-exonic transcripts of multi-exonic genes without known mono-exonic transcripts. Only transcripts observed more than twice across any two samples were retained in the final database of detected transcripts. For the targeted dataset, isoforms associated with only the target genes were retained for downstream analysis.

### Long-read proteogenomic prediction

We used an established long-read proteogenomics pipeline^4^ to generate a comprehensive protein database for identified transcripts. The predicted set of protein isoforms were inferred from identifying different open reading frames (ORFs) using CPAT (v3.0.5)^5^ using all default parameters; transcripts were predicted as protein-coding if the coding potential score was >=0.364. Transcripts that were predicted to produce identical proteins were collapsed into one entry with the most abundant transcript used as the representative RNA isoform. SQANTI3 filters out proteins that are not FSM, NIC, NNC, ISM (either N- or C-terminus truncations) and NNC with junctions after the stop codon (default 2).

### Proteomic analysis of novel isoforms using liquid chromatography-tandem mass spectrometry

An independent set of postnatal human cortex brain samples (n = 35) from the Brains for Dementia Research (BDR) were processed and prepared for label free liquid chromatography-mass spectrometry (LC/MS). The protein isolation process from brain tissue for LC-MS analysis involved tissue grinding under liquid nitrogen, followed by dissolution in urea. After freeze-thaw cycles and thorough vortexing, centrifugation at 14000 rpm for 30 minutes at 10°C yielded a protein-enriched supernatant. Protein concentration was assessed with a Bradford assay (Biorad, Lunteren, The Netherlands). Each sample was loaded onto a 12% SDS-PAGE gel (Biorad), electrophoresed at 180 V for 4 min, and stained with Coomassie blue (Sigma-Aldrich). Cysteines were reduced and alkylated, followed by washing and dehydration. Trypsin (Promega, Leiden, The Netherlands) was added to gel plugs for digestion. Peptides were extracted in triplicate using 1% formic acid (Biosolve) and 2% acetonitrile, followed by volume reduction in a speedvac (Eppendorf, Nijmegen, The Netherlands) to reach a final volume of 50 μL. For LC-MS/MS analysis, proteins were injected, with randomization and inter-run blanks to minimise carryover. Samples were run consecutively to avoid batch effects. Peptides were separated by Acclaim PepMap C18 analytical column (2 μm, 75 μm × 500 mm, 100 Å) using Thermo Fisher Scientific Dionex Ultimate 3000 Rapid Separation ultrahigh-performance liquid-chromatography (HPLC) system (Thermo Scientific, Waltham, MA, USA). Raw proteomics data (raw Thermo format) was compared to a protein database derived from the CPAT ORF predictions on the human cortex transcriptome dataset. Standard proteomic analysis of the tryptic and multi-protease datasets was performed using the open-source search software program MetaMorpheus (v0.0.316)^6^. All spectra files were first converted to MzML format with MSConvert (centroid mode) prior to analysis with MetaMorpheus. All peptide results reported employ a 1% False Discovery Rate (FDR) threshold after target-decoy searching. The output results tables were analyzed using an established long-read proteogenomics pipeline^7^ to determine if each identified peptide was present in the GENCODE reference or was novel.

### Characterisation of alternative splicing (AS) events and transcript visualisation

Common alternative splicing events including alternative first exon use (AF), alternative last exon use (AL), alternative use of 5’ splice sites (A5), alternative use of 3’ splice sites (A3), intron retention (IR) and exon skipping (ES) were assessed. We used *FICLE* (Full Isoform Characterisation from Long-read sequencing Experiments)^4^ to accurately characterize AS events in known protein-coding genes (n = 19,445) detected in our cortex data. The frequency of AS events in prenatal and postnatal cortex were then normalized for each isoform by multiplying the number of observed events with the normalized isoform expression in prenatal and postnatal samples, and then dividing by the total number of observed events across all processed genes. Transcripts were visualised using either the UCSC genome browser or *ggtranscript* (https://github.com/dzhang32/ggtranscript)^5^.

### Differential transcript expression and differential transcript usage analyses

*DESeq2* (v1.42.0) (https://github.com/thelovelab/DESeq2) was used to perform differential expression analysis for transcript- and gene-level read counts^6^. Gene-level expression was calculated by summing all full-length read counts from associated isoforms. Postnatal samples aged less than 1 year old were excluded from differential expression analysis (N=3) given their proximity to birth. Datasets were filtered for single-exon intergenic transcripts, single-exon transcripts from multi-exonic genes and lowly expressed isoforms (minimum of 10 reads across all samples). Sex and age effects were calculated using the Wald test.

Differential transcript use (DTU) analysis was performed using EdgeR *splicevariant* (https://rdrr.io/bioc/edgeR/man/spliceVariants.html). Lowly-expressed isoforms were filtered, whereby an isoform was only retained if its relative proportion to the major isoform was above a specific fold change (FC) threshold (default FC=0.5). The isoform usage at an isoform and sample level was determined by dividing the normalized read count for each associated isoform by the total normalized read counts of all associated isoforms. The isoform usage at an isoform level only was determined by dividing the mean normalized read count for each associated isoform across all the samples by the total normalized read counts of all associated isoforms.

### Conservation of novel coding sequences

Novel coding sequences (CDS) were determined by overlapping the CPAT^5^ predicted coding sequences, from all isoforms in the data, with those from all GENCODE v44^8^ predicted coding sequences. The UTR regions were determined from the GENCODE v44 protein-coding transcripts. For the random intronic and intergenic regions, a random sample of 10,000 base pairs was taken when the regions were larger than 10kb. Each base-pair was only counted once in the case of overlapping transcripts/regions. PhyloP 241-way conservation scores^9^ were then extracted for each base pair of the novel CDS, GENCODE UTR exons, random intronic regions, and random intergenic regions. Highly conserved bases were defined as those with phyloP>2.27, consistent with Sullivan *et al*.^9^. Confidence intervals for the proportion of highly conserved bases in each category were calculated using 50 iterations of bootstrapping. For the disease gene sets, the DDG2P list^10^ (downloaded 26/09/2023; https://www.deciphergenomics.org/) was used for developmental disorders, the SFARI category 1 genes (downloaded 17/07/2023) were used for autism^11^, and the SCHEMA gene list was used for Schizophrenia^12^.

### Integration with genome sequencing data

Exome-sequencing data from schizophrenia patients, bipolar disorder patients and controls were obtained as described previously^12–14^ and variants overlapping identified novel coding sequences were extracted. Re-annotating these to the relevant transcript, we kept variants annotated as missense or more severe, and also synonymous variants as a control. We then used a Fisher’s exact test to estimate the enrichment in cases versus controls. For developmental disorders we restricted our search to genes on the DDG2P list^10^. We removed biallelic genes and those in the confidence category “limited”, resulting in 1,011 genes tested. In the Genomic England 100,000 genomes cohort^15^, *de novo* variants were called and filtered using their custom pipeline. We overlapped the *de novo* variants in the “*stringent_filter”* set (as of February 2024) with the novel CDS in the 1,011 DDG2P genes.

Using the Variant Effect Predictor (VEP)^16^, we annotated the variants to the transcripts with the novel CDS using the custom annotation function. For the predicted non-synonymous variants, we compared the phenotypes of the patients with the known phenotypes of cases of the corresponding single-gene disorder and annotated the variant with gnomADv4 allele frequencies^17^ to ensure that the variant is not common in any population. DisoPred3^18^ was used to determine the function of the end of the TBR1 protein which contained a novel stop-gained variant.

