## Supplementary Figures for "An atlas of expressed transcripts in the prenatal and postnatal human cortex"

**Supplementary Figures - Overview**


| | |
|---|---|
| Figure S1 | Rarefaction curves |
| Figure S2 | Comparison to human cortex PacBio long read dataset |
| Figure S3 | Top 10 genes with highest number of isoforms |
| Figure S4 | Most abundantly expressed novel transcripts annotate to RPS27A |
| Figure S5 | Comparison of prenatal and postnatal transcripts |
| Figure S6 | Transcript diversity between prenatal and postnatal samples |
| Figure S7 | Frequency of alternative splicing events using *FICLE* |
| Figure S8 | Top-ranked known differentially expressed transcript across development |
| Figure S9 | Number of genes characterised with DTE, DTU and DGE |
| Figure S10 | Differential expression of autosomal transcripts between females and males |
| Figure S11 | Differentially expressed transcripts associated with sex |
| Figure S12 | Top-ranked autosomal sex differentially-expressed transcript |
| Figure S13 | Differentially-expressed transcripts associated with sex and development |
| Figure S14 | Differential transcript usage between male and female cortex |
| Figure S15 | CDS and UTR length differences across development |
| Figure S16 | Transcript diversity between whole and targeted transcriptome datasets |

**Supplementary Figure 1: Saturation is reached across our ONT whole transcriptome dataset at the isoform level.**

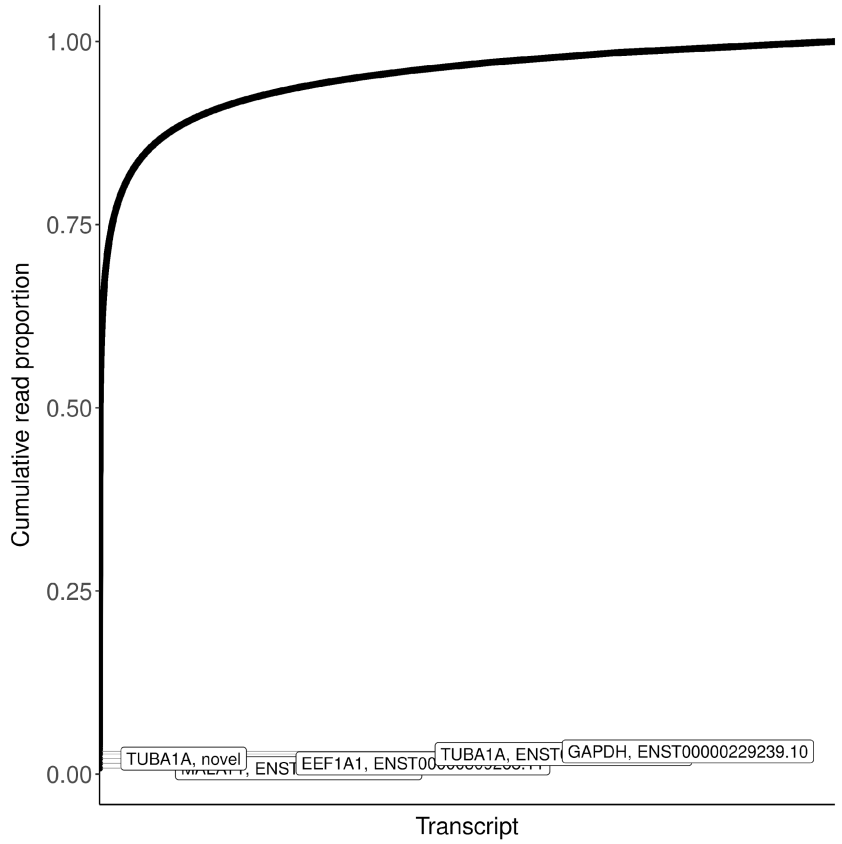

**Supplementary Figure 2: The majority of transcripts identified in our smaller previous analysis of human cortex tissue were detected in the current analysis.** Our previous study used the Pacific Biosciences Iso-Seq approach on a small number of human cortex samples (Leung et al. 2021). The majority of previously detected transcripts (n = 56,252, 86%) were also detected in the current whole transcriptome analysis, with common transcripts identified using the class code "=" in the *GffCompare* package (Pertea and Pertea 2020) (see **Methods**).

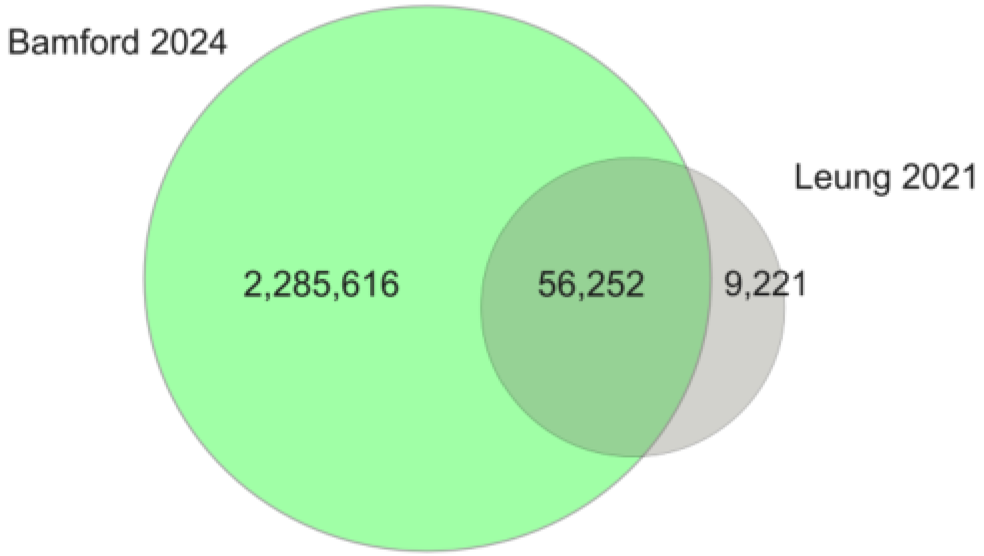

**Supplementary Figure 3: The ten most isomorphic genes identified in the human cortex.** Shown in the left panel is the number of unique transcripts, split by structural category, for each of the ten most isomorphic genes. Shown in the right panel is the number of full-length reads aligning to transcripts of these genes, split by structural category. A large proportion of transcripts annotated to each of these genes are novel (blue and purple), although novel transcripts are individually rare and cumulatively make up only a small proportion of reads.

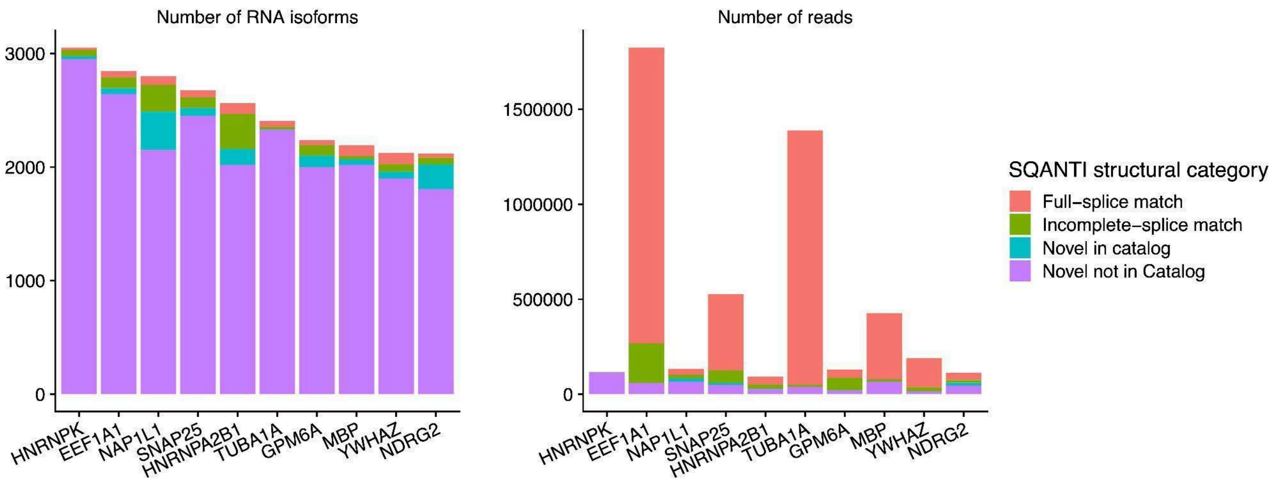

**Supplementary Figure 4: The two most abundantly expressed novel transcripts identified in the human cortex were annotated to the ribosomal protein gene *RPS27A*.** The NNC transcripts ONT2_3223_23931 and ONT2_3223_23974 comprise ~92% of expressed *RPS27A* transcripts. The lines in the bar-plot distinguish the abundance of all distinct transcripts of *RSP27A.*

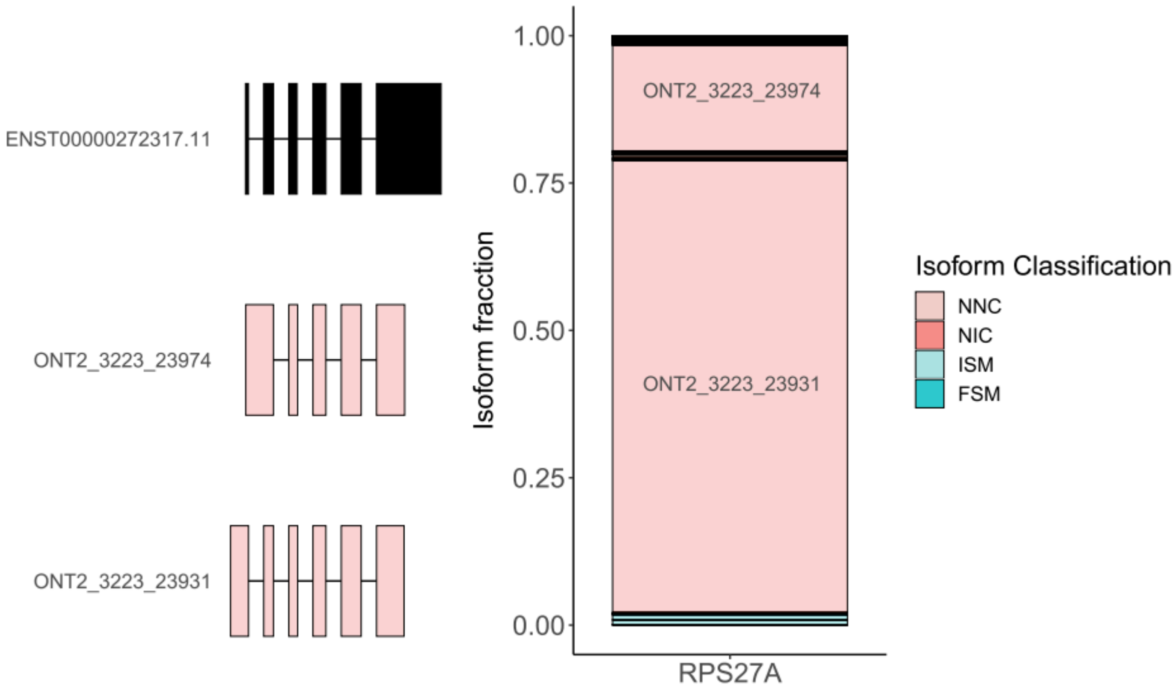

**Supplementary Figure 5: Differences in transcript repertoire between prenatal and postnatal cortex.** Although the majority of transcripts annotated to known genic features were detected in both the prenatal and postnatal cortex (n = 1,184,402 shared transcripts: 75.5% of all detected prenatal transcripts, 78.3% of all detected postnatal transcripts), a large number of transcripts were unique to either the prenatal (n = 383,335; 20% of all detected transcripts) or postnatal (n = 327,623; 17% of all detected transcripts) cortex (see also **Supplementary Table 3**).

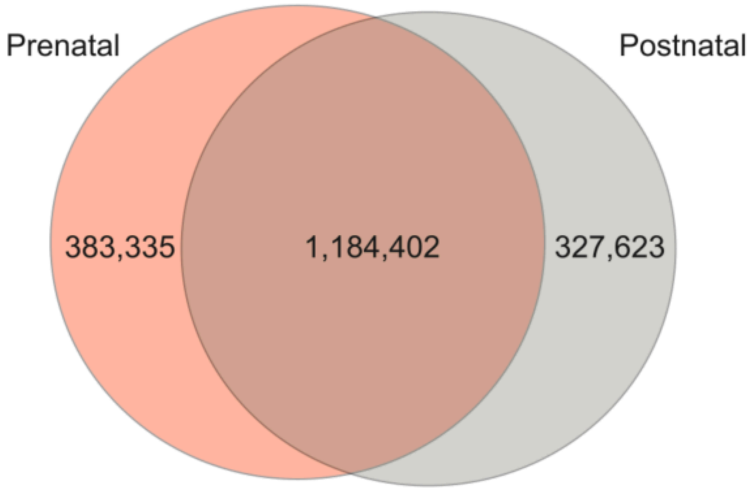

**Supplementary Figure 6: The number of transcripts assigned to different structural categories in prenatal and postnatal cortex.**

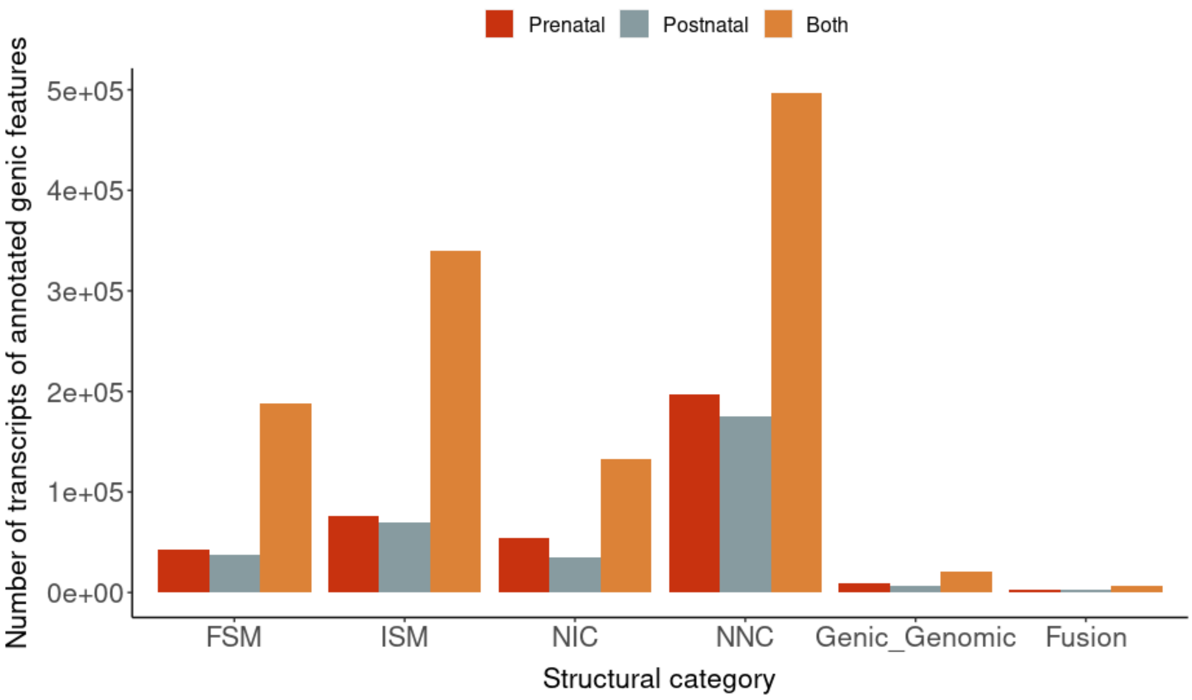

**Supplementary Figure 7: The prevalence of different alternative splicing events in transcripts expressed in the human cortex.** *FICLE* was used to characterise the frequency of different AS events (exon skipping (ES), alternative first exon use (AF), alternative last exon use (AL), alternative A3' and A5' splice site use, and intron retention (IR)) associated with transcripts expressed from multi-exonic protein-coding genes in the human cortex.

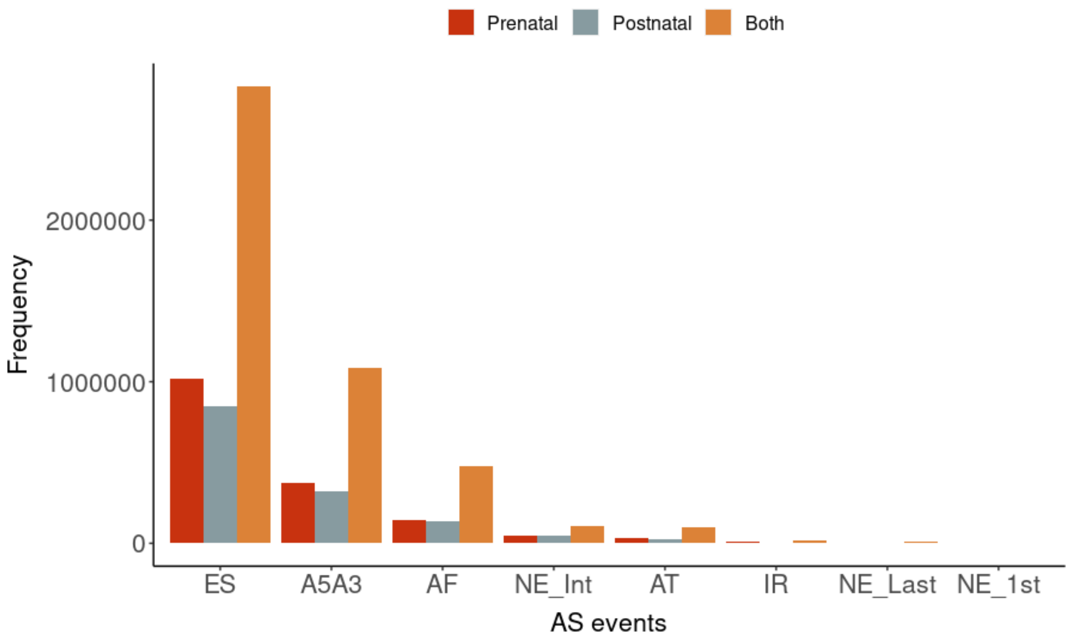

**Supplementary Figure 8: Top-ranked known DET between prenatal and postnatal cortex is an isoform of *MBP*.** Shown is the structure of transcript ONT18_5132_2313 relative to the GENCODEv44 reference transcript and the expression of this transcript in both prenatal and postnatal cortex samples (log2FC = -14.9, FDR = 2.91 x $10^{-142}$).

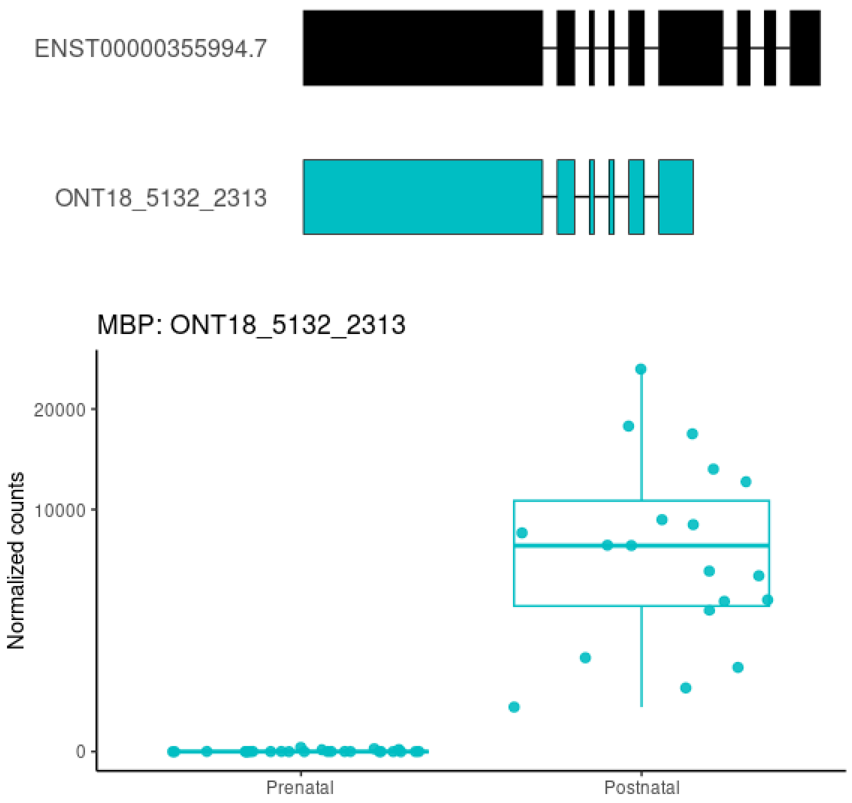

**Supplementary Figure 9: Differential transcript usage and dominant isoform switches between prenatal and postnatal cortex.** Venn diagram showing the number of protein-coding genes that were characterised by significant differential transcript usage across development (DIU), dominant isoform switches, differential gene expression (DGE) and transcripts identified as differentially expressed across development (DTE).

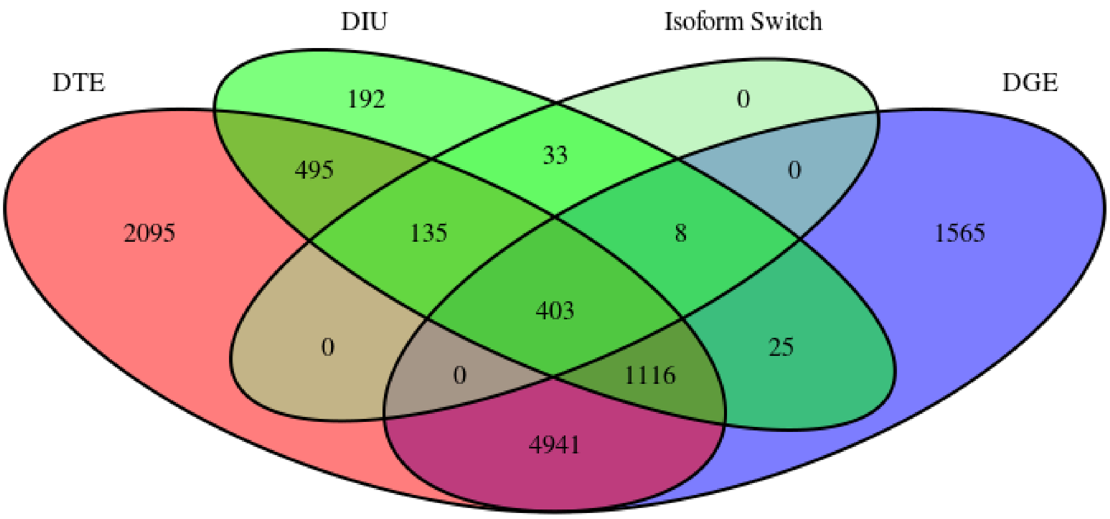

**Supplementary Figure 10: Differential expression of autosomal transcripts between females and males.** Volcano plot highlighting autosomal differentially expressed transcripts (FDR < 0.05) between females and males.

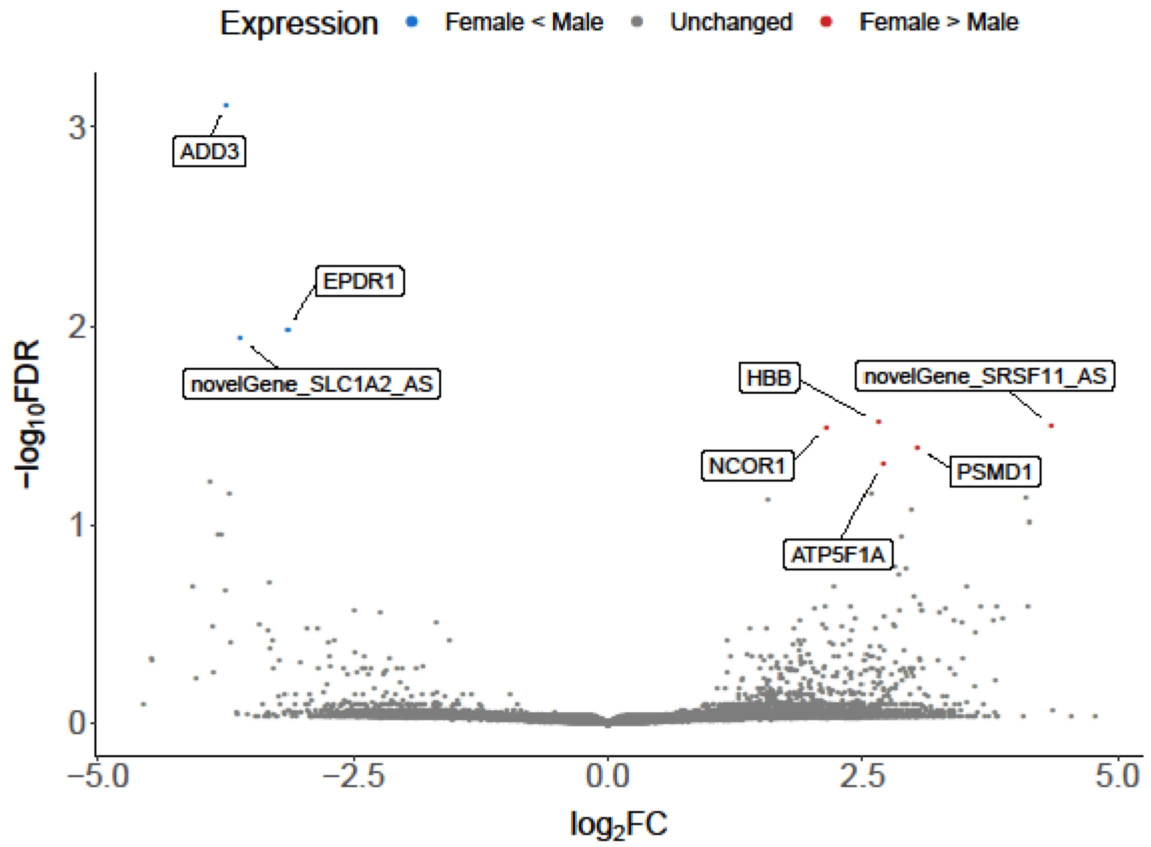

**Supplementary Figure 11: Differential transcript analysis between males and females**. Volcano plot showing the differential expression of 65 transcripts between male and female cortex samples. The 10 top-ranked DETs upregulated in males (blue) and females (red) are highlighted with annotated gene names.

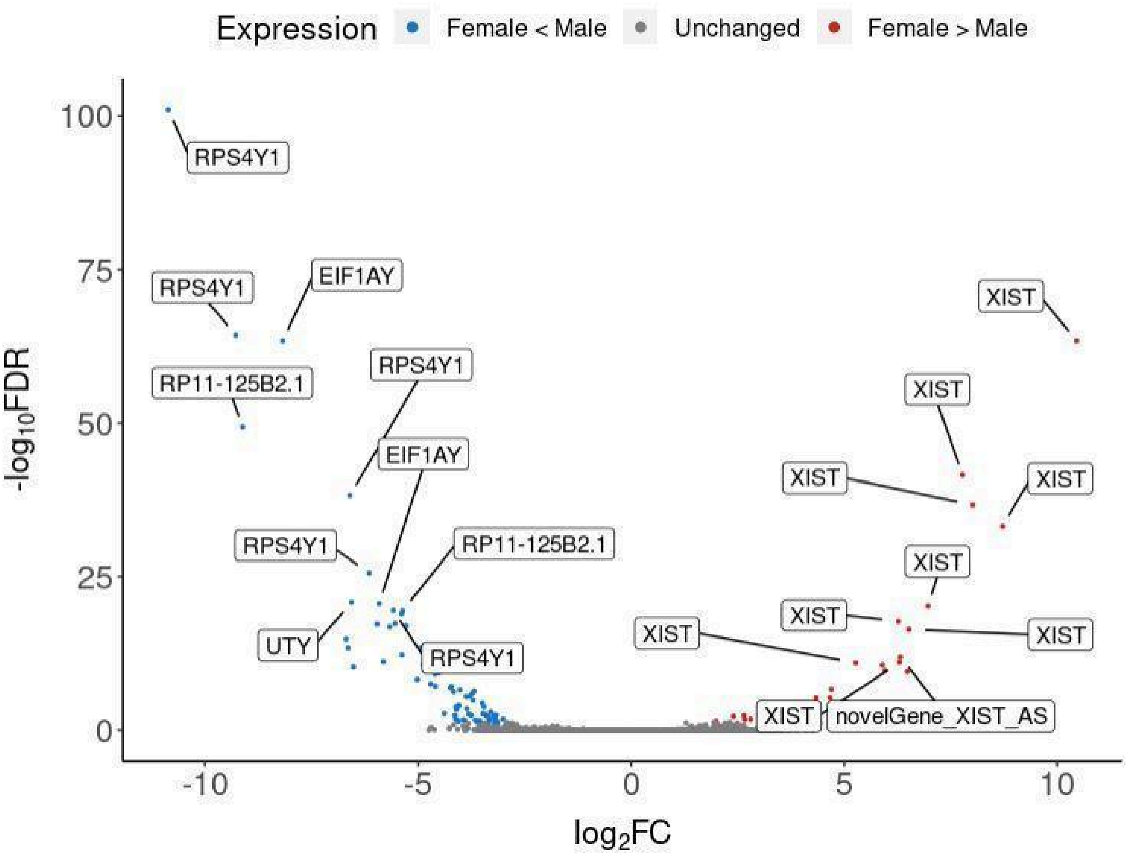

**Supplementary Figure 12: Top-ranked autosomal differentially expressed transcript between males and females.** The top-ranked autosomal sex DET was annotated to *ADD3* (ONT10_4920_1919, log2FC = -3.74, FDR = 7.78 x $10^{-4}$). This transcript was also upregulated in the postnatal cortex only in males (log2FC = -5.21, FDR = 1.60 x $10^{-11}$).

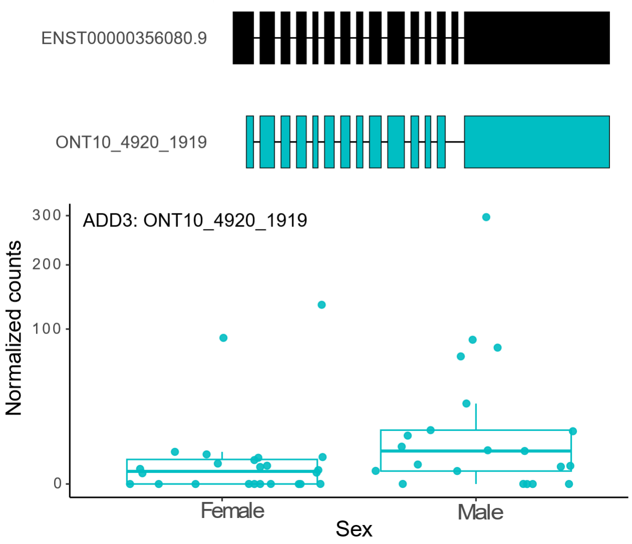

**Supplementary Figure 13: Differential transcript expression across sex and development.** Shown are eight transcripts characterised by both developmental (prenatal vs postnatal) and sex (male vs female) differences in expression . Colours refer to the transcript structural category: blue – FSM, grey – antisense.

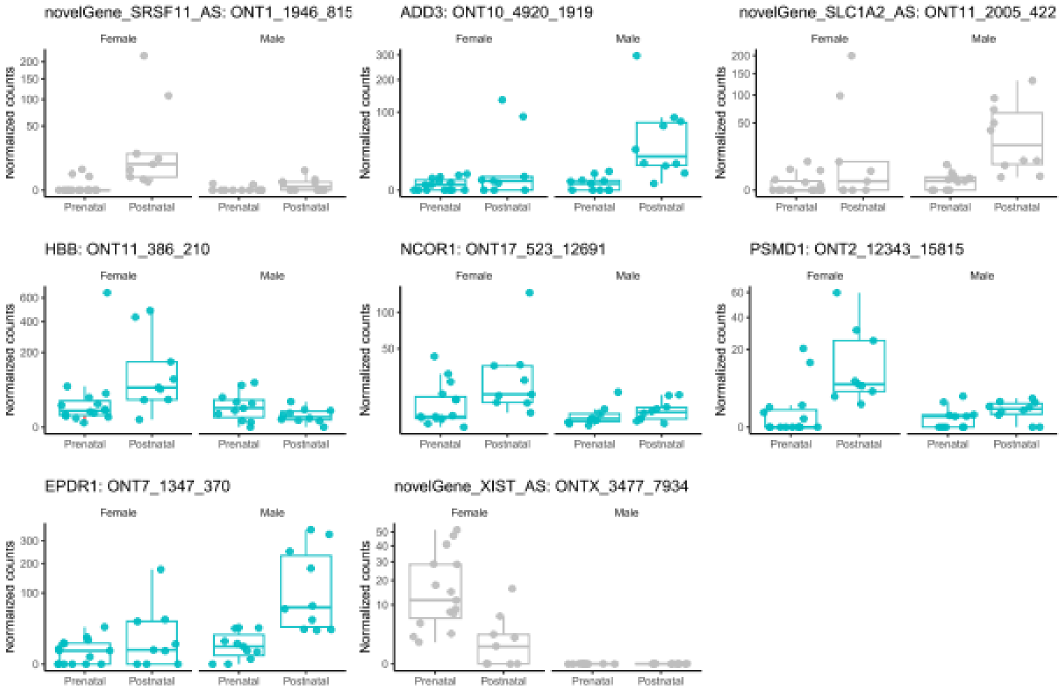

**Supplementary Figure 14: Differential transcript usage between male and female cortex.** Tracks for the top five most abundant transcripts for *GNAS*. Differential transcript use between females and males in the human cortex. Shown for *GNAS* (the top-ranked DTU gene between males and females, p = 1.32 x10$^{-256}$) is the relative isoform fraction for the five most abundant transcripts. There was a switch in major *GNAS* transcript between female and male cortex, with ONT20_3069_10858 being most abundant in female cortex and ONT20_3069_7102 being most abundant in male cortex. There was no overall difference in *GNAS* gene expression between prenatal and postnatal cortex. Tracks are coloured according to structural category (FSM = turquoise; ISM = light turquoise).

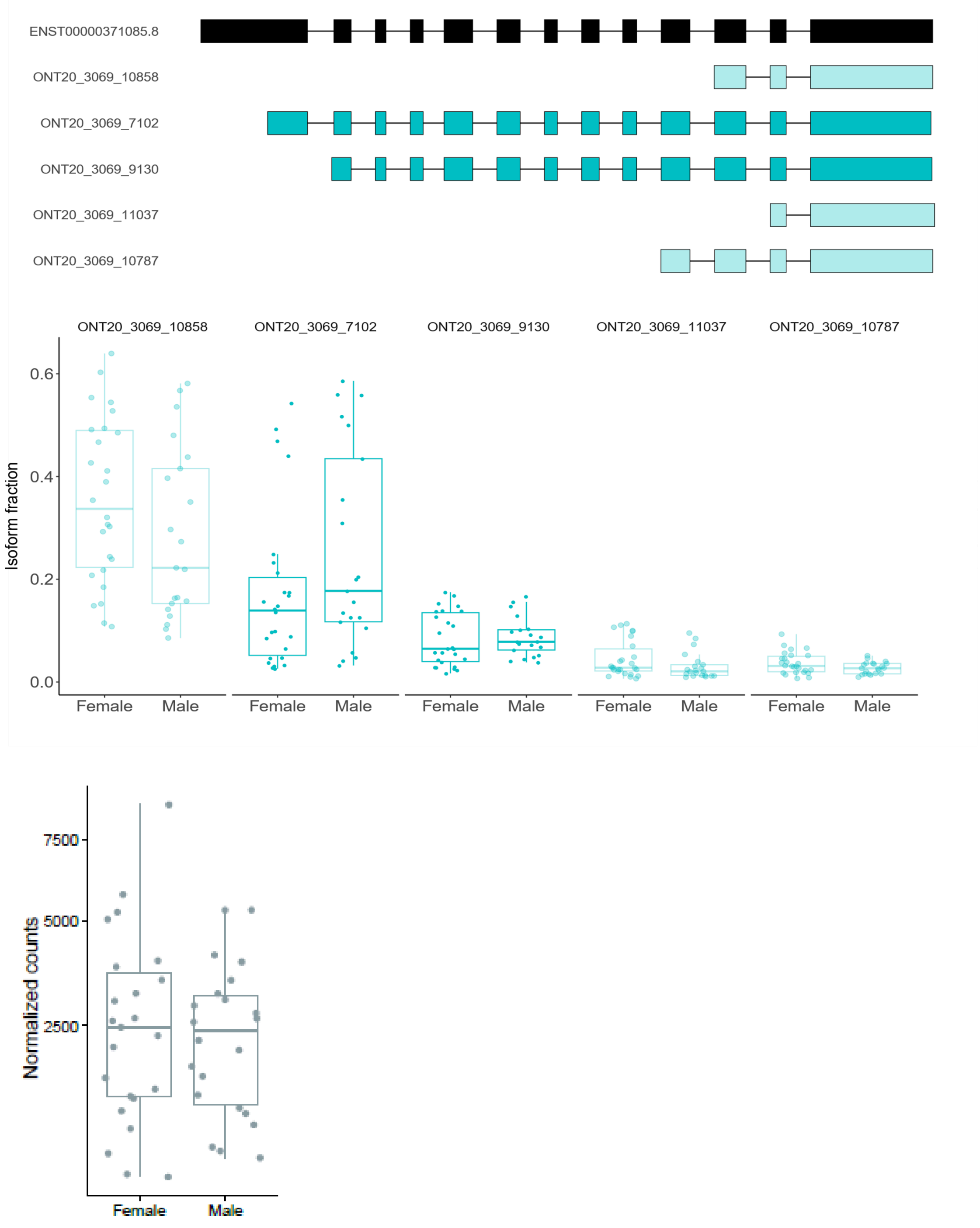

**Supplementary Figure 15: CDS and UTR length differences across development.** The CDS, UTR, 5’ UTR and 3’ UTR length distributions between prenatal and postnatal samples. There is a striking difference between prenatal and postnatal cortex samples in UTR length, which is mainly driven by the 3’ UTR length.

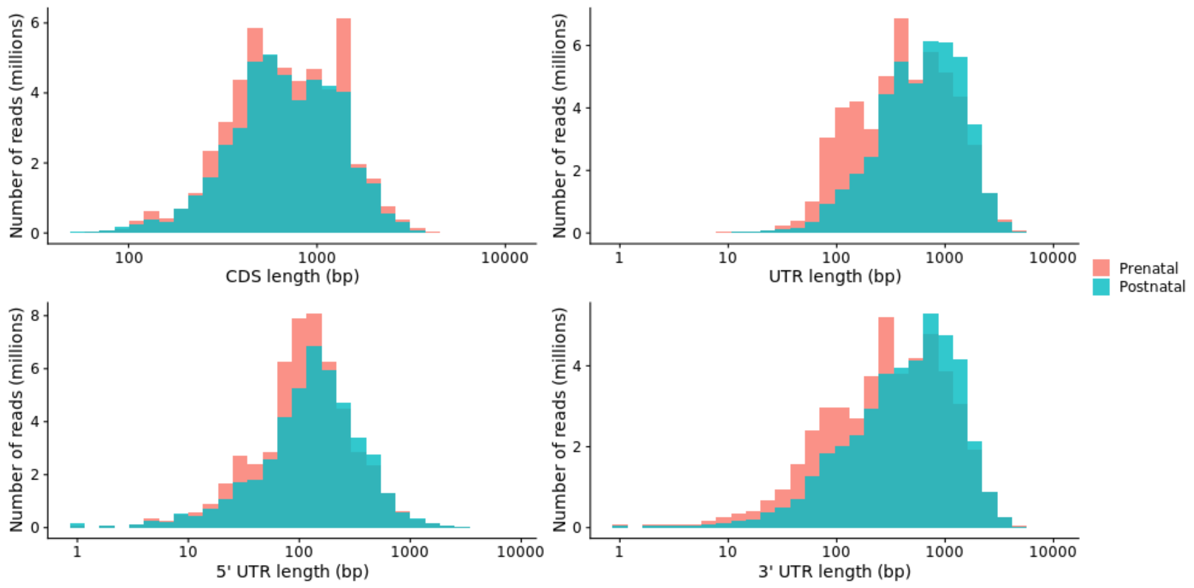

**Supplementary Figure 16: Overlap in isoforms detected using whole and targeted transcriptome sequencing of transcripts from 241 targeted genes.** Shown is **(A)** a Venn diagram of the number of detected isoforms across disease-associated targeted genes in the whole and targeted transcriptome datasets, and **(B)** a bar plot of the number of common and unique of isoforms per disease-associated targeted gene. A full list of genes targeted using this approach is given in **Supplementary Table 15**.

**(A)**

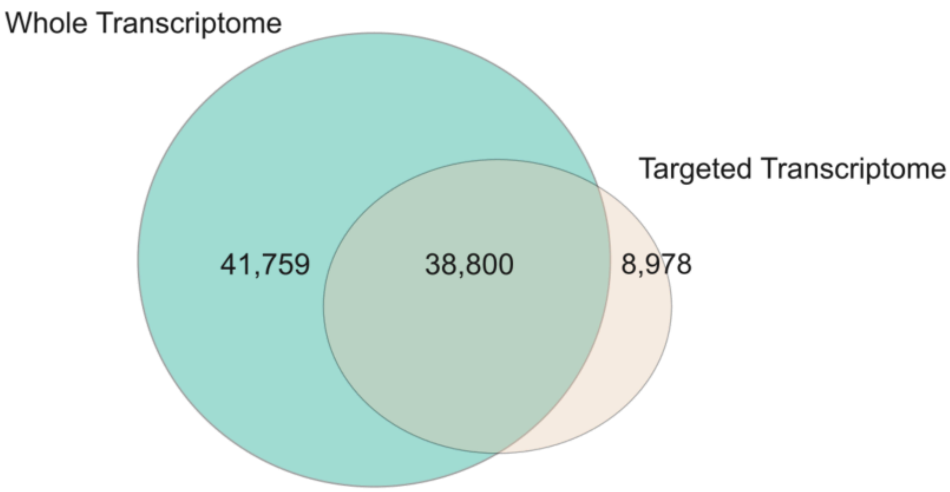


**(B)**

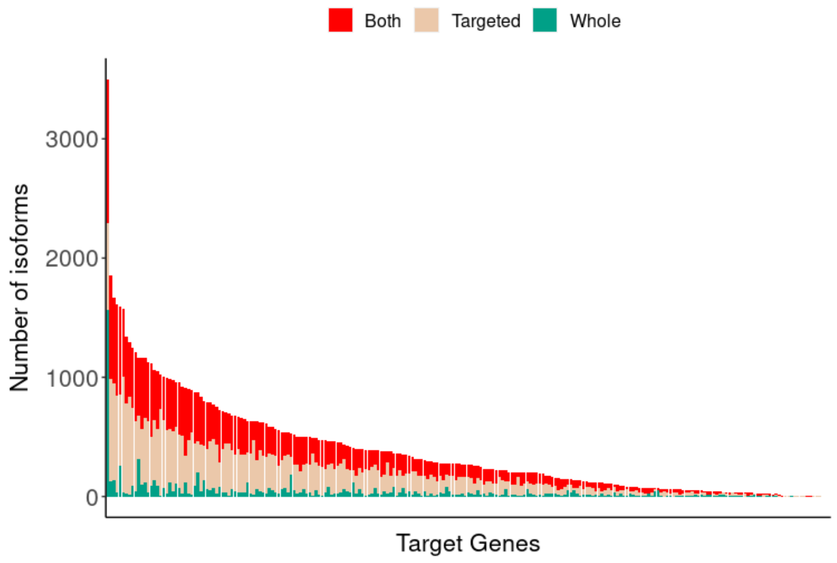
